# DNA methylation shapes the Polycomb landscape during the exit from naïve pluripotency

**DOI:** 10.1101/2023.09.14.557729

**Authors:** Julien Richard Albert, Teresa Urli, Ana Monteagudo-Sánchez, Anna Le Breton, Amina Sultanova, Angélique David, Mathieu Schulz, Maxim V.C. Greenberg

## Abstract

In mammals, 5 methyl-cytosine (5mC) and Polycomb Repressive Complex 2 (PRC2)-deposited histone 3 lysine 27 trimethylation (H3K27me3) are generally mutually exclusive at CpG-rich regions. As mouse embryonic stem cells exit the naïve pluripotent state, there is a massive gain of 5mC coincident with a restriction of broad H3K27me3 to 5mC-free, CpG-rich regions. To formally assess how 5mC shapes the H3K27me3 landscape, we profiled the epigenome of naïve and differentiated cells in the presence and absence of the DNA methylation machinery. Surprisingly, we found that 5mC accumulation is not required to restrict most H3K27me3 domains. We went on to show that this 5mC-independent H3K27me3 restriction is mediated by aberrant expression of the PRC2 antagonist *Ezhip*. At the regions where 5mC appears to genuinely supplant H3K27me3, we identified 68 candidate genes that appeared to require 5mC deposition and/or H3K27me3 depletion for their activation in differentiated cells. Employing site-directed epigenome editing to directly modulate 5mC levels, we demonstrated that 5mC deposition is sufficient to antagonize H3K27me3 deposition and confer gene activation at individual candidates. Altogether, we systematically measured the antagonistic interplay between 5mC and H3K27me3 in a system that recapitulates early embryonic dynamics. Our results suggest that H3K27me3 restraint depends on 5mC, both directly and indirectly. This study also reveals a non-canonical role of 5mC in gene activation, which may be important not only for normal development but also for cancer progression, as oncogenic cells frequently exhibit dynamic replacement of 5mC for H3K27me3 and vice versa.

## Introduction

5-cytosine methylation (5mC) is a highly conserved chemical modification that occurs mainly in the CpG dinucleotide context in vertebrates. 5mC is part of a hierarchical system that impacts gene regulation and can impart epigenetic memory of transcriptional states across cell division. For example, methylated CpG-rich gene promoters are associated with stable transcriptional silencing^1,2^. In this regard, 5mC exhibits properties similar to those of histone 3 lysine 27 trimethylation (H3K27me3), which is deposited by Polycomb Repressive Complex 2 (PRC2) and is also associated with stable silencing when enriched at CpG-rich regulatory regions^3–6^. Given the predominance of methyl-cytosines occurring in the CpG dinucleotide context, we use “5mC” as a signifier of CpG methylation throughout.

In mammals, the fundamental importance of 5mC and H3K27me3 is highlighted by embryonic lethality in mouse knockout studies^7–11^ as well as grossly perturbed 5mC and/or H3K27me3 landscapes in a number of cancers^12–14^. Curiously, mutations in both 5mC and PRC2 machinery are also associated with human overgrowth syndromes^15–19^. The shared ontogenic manifestations of 5mC- and H3K27me3-linked syndromes raise questions about the interplay between the two forms of transcriptional control.

Notably, 5mC and H3K27me3 are largely mutually exclusive at CpG-rich regions (CpG islands, CGIs)—regulatory sequences that frequently overlap gene promoters^20–22^. Components of PRC2^23,24^ and its sister complex PRC1^25^ have been demonstrated to be sensitive to 5mC^26^. Consistently, genetic ablation of DNA methyltransferase encoding genes *Dnmt1, Dnmt3a* and *Dnmt3b* (triple knockout, TKO) in mouse embryonic stem cells (mESCs) results in a complete loss of 5mC and coincident spreading of H3K27me3^21,27^. Conversely, genetic ablation of PRC2 activity results in aberrant 5mC deposition at otherwise H3K27me3-marked regions in mESCs^28^. Hence, Polycomb machinery appears both sensitive to the presence of 5mC, and refractory to DNA methylation deposition. Nevertheless, while the vast majority of CGI promoters are DNA hypomethylated^29^, most of the mammalian genome contains diminished CpG density, and 5mC in this context is the default state. In regions with reduced CpG content, 5mC appears less anticorrelated with H3K27me3^21,22^. Thus, 5mC and H3K27me3 mutual antagonism appears contingent on a threshold of local CpG content.

H3K27me3-5mC reciprocal dynamics is a hallmark of mouse embryogenesis. While pre-implantation blastocysts are DNA hypomethylated and contain broad H3K27me3 domains, genome-wide deposition of 5mC during post-implantation development is coincident with restriction of H3K27me3 to hypomethylated CGIs by the epiblast stage^30–32^. This switch from widespread H3K27me3 to widespread 5mC is of particular interest for developmental progression, as the chromatin established in the epiblast arises just prior to lineage specification, and the epigenetic properties of the marks have the latent potential to confer effects well beyond the precocious embryo. Indeed, aberrant 5mC deposition during epiblast formation can have life-long consequences^33^. However, testing the relevance of H3K27me3 restriction at this stage is complicated by the lethality of embryos lacking 5mC.

The aforementioned H3K27me3-5mC embryonic dynamics occur during the exit from naïve pluripotency, which make pluripotent mESCs an attractive model to dissect the interaction between the two forms of regulation. Past studies have largely relied on metastable mESCs cultured in fetal bovine serum (FBS), which exhibit ∼70% global CpG methylation^21,27,34,35^, and are resilient to genetic ablation of both DNA methylation^36^ and PRC2 machinery^22,37^. However, metastable mESCs represent a heterogenous population of cells consisting of the spectrum of pluripotency states, including “naïve”, “primed”, and the intermediate “formative” state^38,39^. We therefore sought to measure the interplay between 5mC and H3K27me3 in a differentiation system that recapitulates the exit from naïve pluripotency while mitigating cellular heterogeneity in a time-resolved manner^40–42^. To do so, we employed a naïve ESC to epiblast-like cell (EpiLC) differentiation system in which cells pass from a DNA hypo- to hypermethylated state within four days, and exhibit a restricted H3K27me3 landscape within seven^33,43^. Importantly, we previously demonstrated that cells lacking 5mC *(i*.*e*., TKO) successfully exit the naïve state and exhibit hallmarks of primed pluripotent cells, despite substantial transcriptional misregulation^44^. Thus, we used this model to formally assess the effect of genome-wide 5mC deposition on H3K27me3 patterning, and the transcriptional impact on underlying genes.

Unexpectedly, we observed extensive H3K27me3 restriction in TKO EpiLCs despite the complete lack of global 5mC deposition. We showed that in the absence of 5mC, ectopic expression of *Ezhip*, a PRC2 antagonist that is normally silenced in a 5mC-dependent manner, restrains H3K27me3 spreading. A distinct set of loci harbouring intermediate CpG density aberrantly maintained H3K27me3 in TKO EpiLCs, suggesting that 5mC normally supplants H3K27me3 at thousands of regions. Interestingly, a subset of H3K27me3-to-5mC switch regions is associated with transcriptional activation in WT EpiLCs. In other words, genes exhibiting a promoter-proximal H3K27me3-to-5mC switch in WT EpiLCs maintain H3K27me3 enrichment in TKO EpiLCs and fail to properly activate. Single-locus modulation of 5mC levels in naïve and differentiated cells using epigenome editing demonstrated that 5mC can directly antagonize H3K27me3-mediated repression. Previously, we showed in the case of the *Zdbf2* gene not only that embryonic H3K27me3-to-5mC switching is required for activation of this gene, but that failure of this switch results in a life-long growth defect. Here we show that this form of embryonic epigenetic regulation is of a much broader scope and that may potentially impact myriad downstream developmental processes.

## Results

### The epigenome switches from broad H3K27me3 to broad 5mC during the exit from naïve pluripotency

To investigate the interplay between 5mC and PRC2, we performed Whole Genome Bisulfite Sequencing (WGBS) and H3K27me3 Cleavage Under Targets and Tagmentation (CUT&Tag)^45^ in ESCs cultured with double inhibition of ERK and GSK3β supplemented with vitamin C (2i+vitC)^46^ and EpiLCs^40^—a cellular differentiation system that recapitulates the dramatic transition from a globally hypo- to hyper-cytosine methylated genome that occurs during *in vivo* embryonic development (Fig. 1a). Chromatin profiling was complemented with RNA sequencing (RNAseq) in order to correlate DNA and histone methylation dynamics with changes in gene expression. We performed a protracted EpiLC differentiation protocol (7 days) to ensure complete *de novo* deposition of DNA methylation and H3K27me3 turnover.

**Figure 1.**
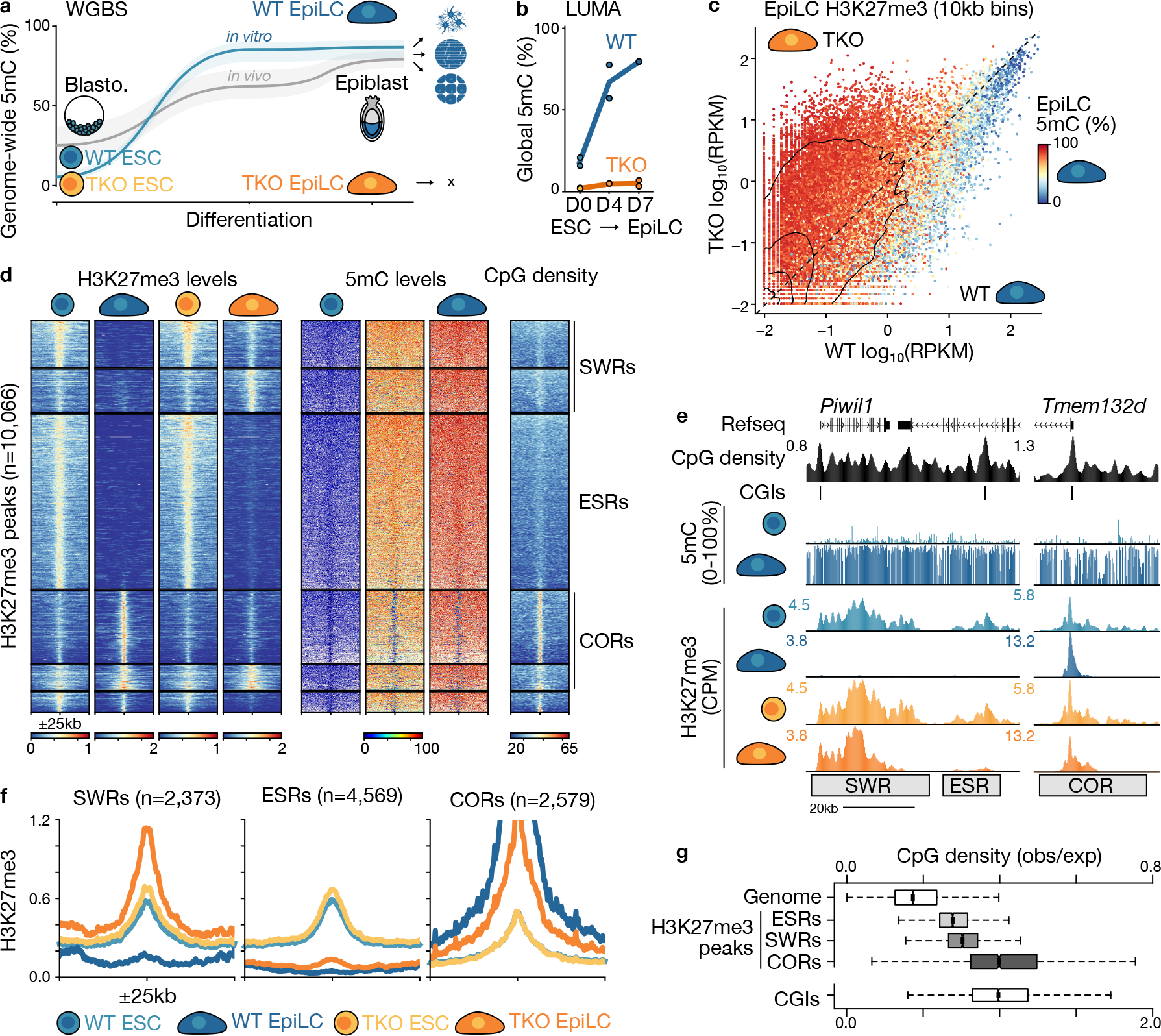
H3K27me3 persists at large fraction of the genome in absence of 5mC. **a.** Whole genome bisulphite sequencing (WGBS) data showing the median (line) and 25^th^ and 75^th^ quartiles (shaded area) of 5mC levels over 10kb bins (n=273,121) during the exit from naïve pluripotency. *In vitro* (blue): 2i+vitC grown ESCs (left), 4-day (middle), and 7-day (right) EpiLCs. *In vivo* (grey): embryonic day 3.5 inner cell mass of the blastocyst (blasto., left), E6.5 (middle) and E7.5 epiblast (right). *In vivo* data from Wang et al. 2014. **b.** Luminometric methylation assay (LUMA) showing global 5mC levels in WT and TKO ESCs and EpiLCs (0, 4 and 7 days post-Activin A/FGF2: AF treatment). **c.** Scatterplot showing global H3K27me3 levels over 10kb bins in WT and TKO EpiLCs (AF treatment day 7). Datapoints are coloured by average 5mC levels in WT EpiLCs. 10kb bins with ≥10 CpGs covered by ≥5 reads are shown (n=252,559); contours are drawn at iso-proportions of the density of data points. **d.** Heat map of k-means clustered H3K27me3 peaks, showing H3K27me3, 5mC and CpG density levels 25kb up- and downstream peak centers (n=10,066). Three groups of H3K27me3 peaks are indicated: H3K27me3-to-5mC Switch Region (SWR), ESC-Specific Region (ESR) and Constitutive Region (COR) **e.** UCSC Genome Browser screenshot of an example SWR, ESR and COR. Data from WT and TKO ESC and EpiLCs (AF treatment day 7) is shown. CPM: counts per million aligned reads. Coordinates: chr5:128,725,246-128,800,866 (left) and chr5:128,399,926-128,475,639 (right). **f.** Metaplot showing H3K27me3 enrichment over three H3K27me3 peak categories defined in **d** in WT and TKO ESCs and EpiLCs. **g.** Boxplot representing the distribution of CpG density levels genome-wide, over CpG islands (CGIs) and over H3K27me3 peaks (SWRs, ESRs and CORs). Note: CGIs are placed on a different scale.

During differentiation, 5mC and H3K27me3 levels over CpG islands were largely static (Fig. S1a). That is to say, the CGI promoters of PRC2 target genes remained hypomethylated and maintained high levels of H3K27me3 in EpiLCs (Fig. S1a-b) in association with transcriptional silencing (Fig. S1c). Only a minority of CGI promoters exhibited dynamic H3K27me3 patterns that generally anticorrelated with the transcriptional state. For example, genes active in EpiLCs showed depletion of H3K27me3 over the course of differentiation (Fig. S1a,d,e). Conversely, naïve pluripotency marker genes gain H3K27me3 in both EpiLCs and the epiblast (fig. S1f,g). Finally, a small subset of CGIs—including the promoters of germline-expressed genes that are known to be DNA methylation-regulated—gain 5mC (>75%) at the expense of H3K27me3 in EpiLCs (Fig. S2a). This switch from H3K27me3 to dense 5mC levels was associated with transcriptional silencing in both cell types, with the notable feature that at least some germline genes exhibited a transient transcriptional burst during the H3K27me3-5mC transition both *in vitro* and *in vivo* (Fig. S2b). Consistent with our differentiation model, 66% of CGI promoters that significantly (linear modelling using Limma, fold-change >2, t-test adjusted p<0.05) lose H3K27me3 coincident with gain of 5mC in WT EpiLCs also gain 5mC in the developing epiblast (Fig. S2c). Nevertheless, CGIs are generally devoid of DNA methylation (<25% 5mC) in our EpiLC differentiation system (88% of CGIs) and during normal development (92% of CGIs) (Fig. S2d).

Unlike CGIs, the rest of the genome showed widespread H3K27me3 level loss during EpiLC differentiation (Fig. S3a). Notably, 93% of regions that lost H3K27me3 gained >75% 5mC levels in EpiLCs (Fig. S3b), suggesting that 5mC may broadly antagonize H3K27me3 deposition. To test whether 5mC drives H3K27me3 loss during EpiLC differentiation, we performed parallel experiments in TKO cells that are completely devoid of 5mC (Fig. 1b)^36,44,47^. Analysis of genome-wide H3K27me3 levels showed a strong correlation between globally hypomethylated wild-type (WT) and TKO ESCs (Fig. S3c). Notably, relative to WT EpiLCs, TKO EpiLCs exhibited a clearly elevated H3K27me3 distribution (Fig. 1c). Consistent with our observations that 5mC restricts H3K27me3 over CGIs, 94% of regions that aberrantly maintain H3K27me3 in TKO EpiLCs were normally hypermethylated (>75% 5mC) in WT EpiLCs (Fig. S3d). It should be noted that elevated levels of H3K27me3 in TKO EpiLCs could potentially be confounded by differentiation defects; however, aberrantly H3K27me3-marked regions in TKO EpiLCs included less than half of those that have H3K27me3 in ESCs as well as thousands of additional regions, indicating that TKO EpiLCs do not simply exhibit a naïve ESC profile (Fig. S3e). Further supporting this notion, analysis of marker gene expression indicated that differentiation occurred with similar dynamics even in the absence of 5mC, which is consistent with previous reports (Fig. S4a)^44^. Moreover, principal component analysis of WT and TKO RNAseq data with *in vivo* data shows clustering of ESCs with blastocyst-stage, and EpiLCs with epiblast-stage embryos, regardless of genetic background (Fig. S4b). These results suggest that 5mC restricts H3K27me3 levels genome-wide during normal EpiLC differentiation and that, in the absence of 5mC, H3K27me3 can remain relatively broadly distributed.

To quantify H3K27me3-to-5mC dynamics, we performed agnostic k-means clustering of H3K27me3 peaks (Fig. 1d-e). Three primary groups emerged: 1) <u>CO</u>nstitutive <u>R</u>egions (CORs) that largely overlapped with CGIs and showed H3K27me3 enrichment and hypomethylation regardless of cell type; 2) <u>E</u>SC-<u>S</u>pecific <u>R</u>egions (ESRs), which lost H3K27me3 levels in both WT and TKO EpiLCs; and 3) H3K27me3-to-5mC <u>SW</u>itch <u>R</u>egions (SWRs) in WT EpiLCs, which aberrantly maintained H3K27me3 levels in TKO EpiLCs. In other words, the H3K27me3 landscape is shaped in a 5mC-independent (ESR) and dependent (SWR) manner.

CpG islands, which are a general feature of CORs (Fig. S4c), are known to be protected from DNA methylation via a number of mechanisms^48^; thus, we were more intrigued to investigate the features that delineate the other two classes. Both ESRs (n=4,569) and SWRs (n=2,373) exhibited similar H3K27me3 levels in ESCs, indicating that H3K27me3 enrichment prior to differentiation was not a predictor of H3K27me3 or 5mC levels in EpiLCs (Fig. 1f). We next wondered if specific transcription factor binding motifs might explain the differential H3K27me3 dynamics. Given that the H3K27me3 domains are quite large, we focused our analysis on 1kb regions in ESRs and SWRs that overlapped with publicly available Polycomb complex subunit binding data^34^. While SWRs were enriched for GC and CG dinucleotides (Fig. S4d), no clear transcription factor binding motifs were found that would indicate specific regulation for either ESRs or SWRs.

PRC2 in mammals has a well-described affinity for CpG-rich regions, such as CGIs^49^. Consistently, all three classes of H3K27me3-enriched regions exhibited CpG density higher than genome average (Fig. 1d,g). However, ESRs contained the lowest CpG density, with SWRs intermediate, and CORs drastically higher. This indicates that in WT EpiLCs, PRC2 affinity is shifted towards regions with higher CpG content (CORs), and that the gain of DNA methylation specifically at CpG-rich regions (ESRs, SWRs) is refractory to PRC2 activity. Nevertheless, an intriguing question arose: what accounted for the decrease of H3K27me3 deposition not only in WT, but also in TKO EpiLCs at a large fraction of the genome (ESRs)?

### H3K36me3 deposition is unlinked to H3K27me3 misregulation in TKO EpiLCs

Given the H3K27me3 loss concomitant with 5mC gain during the exit from the naïve state, it was somewhat surprising that such a large proportion of regions lost the histone mark in the complete absence of DNA methylation machinery. In fact, ESRs comprised the greatest proportion of H3K27me3-marked regions (Fig. 1d). Concordantly, genome-wide loss of H3K27me3 levels was observed in differentiating TKO cells (Fig. 2a). We hypothesized that perhaps H3K36 methylation, which has been shown to antagonize H3K27me3 *in vitro*^50,51^, may play a role in limiting broad H3K27me3 deposition. Indeed, both *in vivo* and cell culture models of H3K36 misregulation have reported extensive H3K27me3 remodeling^27,52^. Given the link between H3K36me2/3 and 5mC^53–55^, we tested whether the H3K36 methylation landscape is altered in TKO EpiLCs. As expected, H3K36me3 CUT&Tag analysis showed a clear anticorrelation between H3K36me3 and H3K27me3 in ESCs and EpiLCs (Fig. S5a-b), and the vast majority of H3K36me3-marked sites are DNA methylated in WT EpiLCs (Fig. S5b). However, differential analysis in WT versus TKO showed highly correlative H3K36me3 patterns in ESCs and EpiLCs (Fig. S5c-d). Indeed, regions that showed increased H3K27me3 levels in TKO EpiLCs relative to WT EpiLCs did not exhibit a concomitant depletion of H3K36me3 (Fig. S5e-f). While we did not formally exclude a role for H3K36me2 misregulation in TKO EpiLCs, previous reports have demonstrated that H3K36me2 is broadly deposited in mouse pluripotent cells, and not perturbed in DNA methylation mutants^27,56^. As such, we proceeded to search for alternative explanations for the observed restriction of H3K27me3 in TKO EpiLCs.

**Figure 2.**
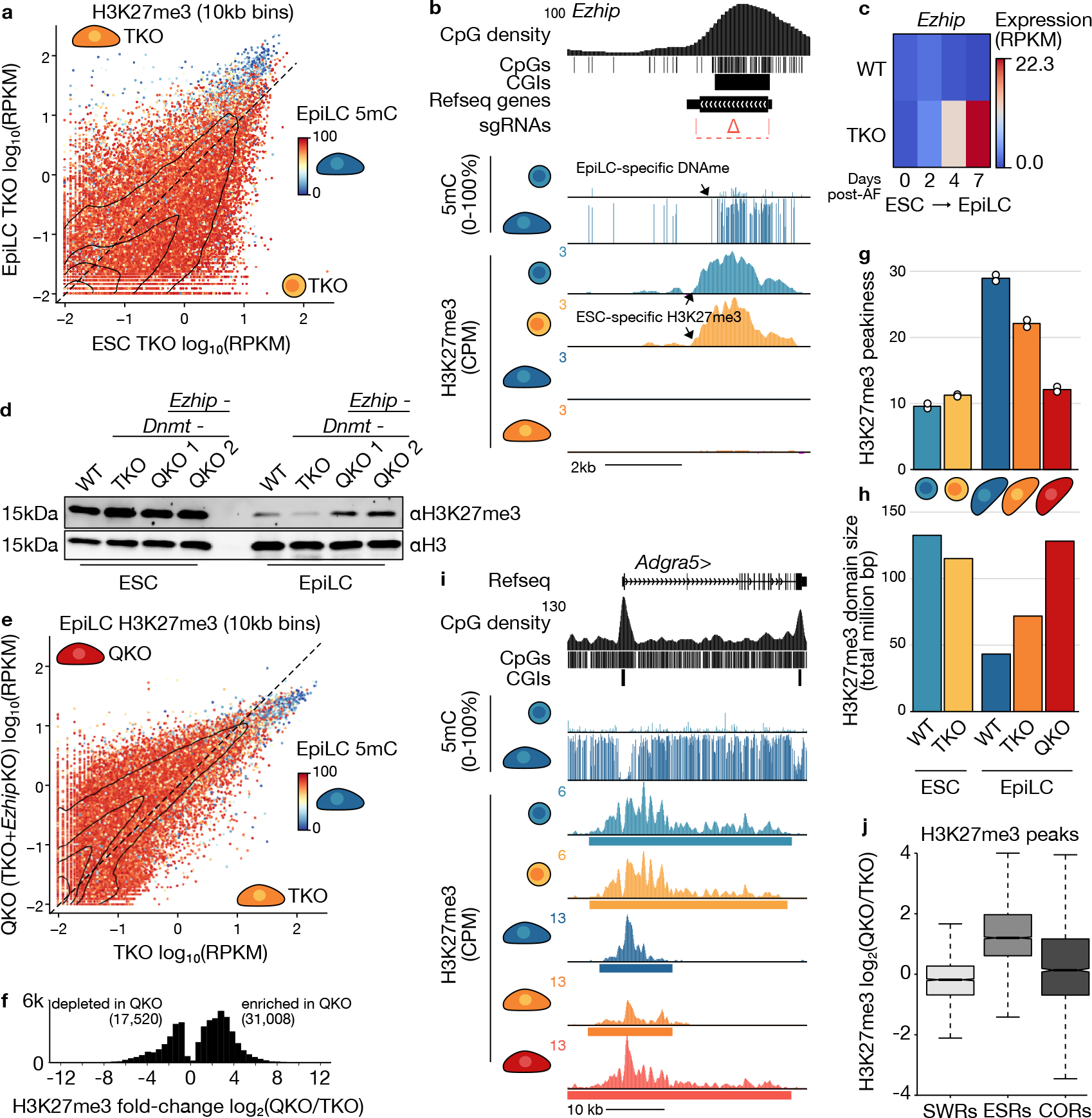
5mC indirectly regulates H3K27me3 spreading via *Ezhip* silencing. **a.** 2D scatterplot showing H3K27me3 enrichment over genome-wide 10kb bins in differentiating TKO cells. Data points are coloured by 5mC levels in WT EpiLCs. 10kb bins with ≥10 CpGs covered by ≥5 reads are shown (n=252,559), and the density of datapoints is included. **b.** UCSC Genome Browser screenshot of the *Ezhip* locus displaying H3K27me3 enrichment over the CGI promoter in WT and TKO ESCs and 5mC enrichment in WT EpiLCs (AF treatment day 4). The location of CRISPR-Cas9 gRNAs and deleted region is shown. Coordinates: chrX:6,078,117-6,084,409. **c.** Heatmap showing *Ezhip* expression levels in WT and TKO ESCs and EpiLCs. EpiLC data was generated from cells 0, 2, 4 and 7 days post-AF treatment. The mean of two replicates is shown. **d.** Western blot showing global H3K27me3 levels in WT, TKO, and two QKO (TKO+*Ezhip* KO) subclones in ESCs and EpiLCs (AF treatment day 7). **e.** 2D scatterplot showing H3K27me3 enrichment over genome-wide 10kb bins in TKO and QKO EpiLCs. Data points are coloured by 5mC levels in WT EpiLCs as in **a.** **f.** Histogram displaying the number of 10kb bins significantly enriched or depleted (defined using linear modeling in Limma, adjusted p value <0.05) for H3K27me3 in QKO EpiLCs relative to TKO EpiLCs. **g.** Bar plot of H3K27me3 domain peakiness scores. Peakiness is defined as the average RPKM level of the top 1% most enriched H3K27me3 peaks. **h.** Bar plot showing the sum of all H3K27me3 peak sizes. Note the anti-correlation with peakiness scores. **i.** UCSC Genome Browser screenshot of the *Adgra5* gene showing 5mC-independent, *Ezhip*-dependent H3K27me3 restriction during EpiLC differentiation. H3K27me3 peak calls are shown as rectangles. Coordinates: chr8:27,074,539-27,123,049. **j.** Boxplot showing the relative change in H3K27me3 level enrichment over H3K27me3 peaks between TKO and QKO EpiLCs.

### Ectopic Ezhip expression mitigates H3K27me3 spreading in TKO EpiLCs

Recently a number of papers have described EZHIP as a PRC2 antagonist in mouse germline development and in some human cancers^57–61^. *Ezhip* was strongly enriched for H3K27me3 in ESCs (Fig. 2b) and published expression data in ESCs lacking PRC2 components exhibit *Ezhip* upregulation (Fig. S6a)^62,63^. In WT EpiLCs, the *Ezhip* promoter exhibited dense 5mC and was transcriptionally repressed (Fig. 2b-c). However, unmethylated *Ezhip* was highly transcribed (∼24 RPKM) in TKO EpiLCs (>300-fold upregulated compared to WT) (Fig. 2c).

To determine if *Ezhip* expression restricts H3K27me3 in EpiLCs lacking 5mC, we used CRISPR-Cas9 to genetically ablate *Ezhip* in TKO ESCs and performed EpiLC differentiation. TKO+*Ezhip* KO (QKO) EpiLCs resembled both WT and TKO EpiLCs morphologically, and expressed marker genes at comparable levels (Fig. S6b-c). Western blot analysis revealed similar global H3K27me3 levels in WT and TKO ESCs and EpiLCs, with a reduction of H3K27me3 signal in the EpiLC state (Fig. 2d). Strikingly, two QKO lines showed a global increase in H3K27me3 levels compared to TKO EpiLCs (Fig. 2d). It should be noted that QKO H3K27me3 levels did not appear as high as in ESCs, which may be due to downregulation of core PRC2 machinery at the RNA and protein level in EpiLCs (Fig. S6d-e)^64^. Nevertheless, genome-wide analysis of H3K27me3 distribution by CUT&Tag in QKO versus TKO EpiLCs uncovered increased enrichment of H3K27me3 in cells lacking *Ezhip*, indicative of spreading (Fig. 2e-f). Indeed, H3K27me3 was most restricted in WT EpiLCs, relatively intermediately spread in TKO EpiLCs, and broadly distributed in QKO EpiLCs (Fig. 2g-h). The broad H3K27me3 profiles of QKO EpiLCs was most reminiscent of ESCs (Fig. 2i), suggesting that in the absence of 5mC, differentiating cells can nevertheless restrict H3K27me3 through *Ezhip* expression. EZHIP was a more effective PRC2 antagonist at ESRs, which exhibit less CpG density, and thus potentially less PRC2 affinity than at CORs and SWRs (Fig. 2j). We conclude from our genetic analyses that H3K27me3 can be directly restricted by global deposition of DNA methylation, and via ectopic *Ezhip* expression in the absence of 5mC.

### 5mC accumulates at a set of genes upregulated during EpiLC differentiation

Despite ESRs comprising the largest class of H3K27me3-marked regions, a sizeable fraction (SWRs, ∼25%) maintained H3K27me3 in TKO EpiLCs, even in the presence of EZHIP (Fig. 1d). Given the relatively higher affinity for PRC2 activity, we were curious to know if SWRs spanned regions wherein H3K27me3 exerts a gene-regulatory effect, and by extension, if DNA methylation deposition impacted PRC2-mediated gene control.

Canonically, 5mC is a mark of transcriptional repression in CpG-rich regulatory regions, such as CGIs. At CGIs, a SWR would presumably not lead to a change in transcriptional activity, since H3K27me3 is supplanted by 5mC (inactivating SWR, Fig. S7a). For example, the germline gene *Piwil1* CGI promoter normally acquires 5mC during EpiLC differentiation, however H3K27me3 was aberrantly maintained in TKO EpiLCs (Fig. S7b), suggesting both pathways converge to silence this gene. Indeed, *Piwil1* is repressed (RPKM<0.5) in WT and TKO ESCs and EpiLCs (Fig. S7c).

While 2,426 genes were upregulated in TKO EpiLCs, intriguingly, a substantial set of 1,048 genes were significantly downregulated in TKO EpiLCs (Fig. 3a). This could either reflect indirect effects of DNA methylation loss or examples where 5mC plays a non-canonical activating role (activating SWR). Giving credence to the latter possibility, our previous work demonstrated that the *Zdbf2* gene requires 5mC to displace H3K27me3 in order to ensure its expression during embryogenesis^33^. However, to our knowledge *Zdbf2* is the only gene to be described as regulated in such a manner during this window of differentiation. To determine whether 5mC deposition contributes to gene activation, we intersected a bolstered list of statistically significant SWRs (see Methods, n=2,418) with genes significantly downregulated in TKO EpiLCs (Fig. 3b). SWRs overlapped the promoters of 68 downregulated genes in TKO EpiLCs, including *Zdbf2* (Fig. 3a-c). Importantly, 37 are downregulated in *Dnmt1* KO epiblasts, including *Zdbf2* and *Celsr2*, which substantiates our *in cellula* model (Fig. 3c)^65^. H3K27me3 enrichment is likely responsible for repression, as the majority (48/56) of candidate SWR-activated genes expressed in ESCs are upregulated upon *Suz12* KO (Fig. 3c)^62^. Additionally, 55/68 candidate SWR genes are upregulated in ESCs harboring a triple knockout for PRC1 subunits *Pcgf3, 5* and *6*, which indicates that both Polycomb complexes cooperate to repress expression of SWR genes in hypomethylated cells (Fig. 3c)^66^.

**Figure 3.**
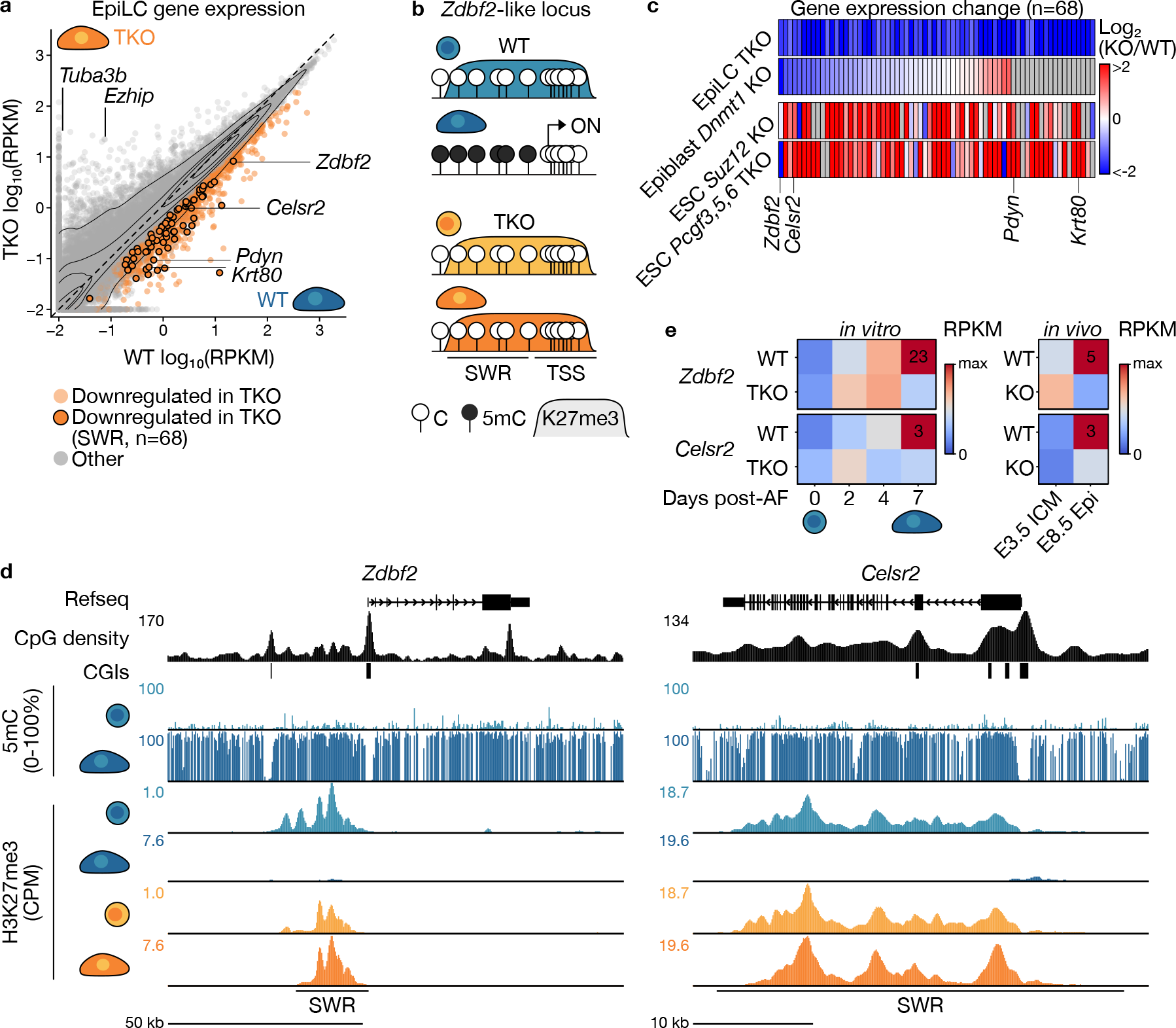
5mC may facilitate the activation of a subset of genes during differentiation. **a.** 2D scatterplot showing gene expression levels in WT and TKO EpiLCs (7 days post-AF). Significantly downregulated genes in TKO EpiLCs (n=1,048) are coloured in orange. Significantly downregulated genes whose transcription start site overlaps a SWR are indicated (n=68), and the density of datapoints is included. **b.** Schema showing H3K27me3 and gene expression patterns used to identify SWR regions that may be sensitive to 5mC-mediated gene activation. **c.** Heatmap showing relative expression values of candidate 5mC-activated genes near SWRs (n=68). Each column displays the fold-change between KO (TKO, *Dnmt1* KO, *Suz12* KO and *Pcgf3,5,6* TKO) and control WT cells in log_2_ scale. Lowly expressed genes in both conditions (average CPM<1) are shown in grey. See Supplemental Table 1 for the full list of data mined in this study. **d.** UCSC Genome Browser screenshots of candidate 5mC-activated genes *Zdbf2* and *Celsr2* adjacent to SWRs. *Zdbf2* chr1:63,222,008-63,338,671; *Celsr2* chr3:108,388,342-108,426,060. **e.** Heatmap displaying *Zdbf2* and *Celsr2* expression levels (RPKM) during differentiation *in vitro* (0, 2, 4 and 7 days post-AF) in WT and TKO cells. Data derived from the developing mouse embryo is included: *Dnmt3a* maternal KO embryonic day 3.5 inner cell mass (ICM) cells and *Dnmt1* zygotic KO embryonic day 8.5 epiblast cells, and control WT embryos. Note the lower expression of *Zdbf2* and *Celsr2* in cells lacking 5mC deposition machinery.

Integrated analysis of 5mC, H3K27me3 and expression levels revealed that broad H3K27me3 upstream and overlapping the *Zdbf2* CGI promoter is normally restricted in WT EpiLCs in association with 5mC deposition and gene activation (Fig. 3d). In contrast, TKO EpiLCs aberrantly maintained H3K27me3, which extends into the CGI promoter and is linked with maintained transcriptional repression (Fig. 3d-e). A similar pattern of 5mC-mediated H3K27me3 restriction and gene activation was observed at the other 67 candidate genes including *Celsr2, Pdyn, Krt80*, and *Pga5* (Fig. 3a-d, Fig. S7d). These data suggest that 5mC may play an activating role in gene expression through Polycomb antagonism at a much larger set of genes than was previously realized as cells differentiate from the naïve state.

### Precision DNA methylation editing at the Zdbf2 SWR is sufficient to antagonise H3K27me3

While our genomic analyses in EpiLCs enabled us to characterize and identify candidate 5mC-activated genes near SWRs, we could not exclude potentially misleading confounding effects. To directly assess the effect of 5mC deposition on H3K27me3 restriction and nearby gene expression, we established and employed two complementary site-directed epigenome editing approaches.

First, we generated an inducible cell line to selectively target 5mC to candidate SWRs in hypomethylated ESCs (Fig. 4a). Briefly, two doxycycline (Dox)-inducible constructs were stably integrated in the two alleles of the *Rosa26* locus, respectively: a catalytically inactive *Cas9* fused to ten GCN4 epitopes (dCas9-SunTag)^67^, and a *Dnmt3a* catalytic domain (3ACD) fused to GFP and a single-chain antibody that recognizes the GCN4 epitope (Fig. S8a). Both constructs contain a FKBP12-derived destabilizing domain, minimizing leaky expression in uninduced cells, and enabling robust expression in the presence of Dox and the Shield-1 ligand (Fig. S8b)^68,69^. Cells expressing a catalytically dead 3ACD (d3ACD) were used as a control. Finally, we stably integrated single-guide RNAs (sgRNAs) designed to target the predicted Polycomb nucleation site (see Methods) of candidate SWRs (Fig. S8c).

**Figure 4.**
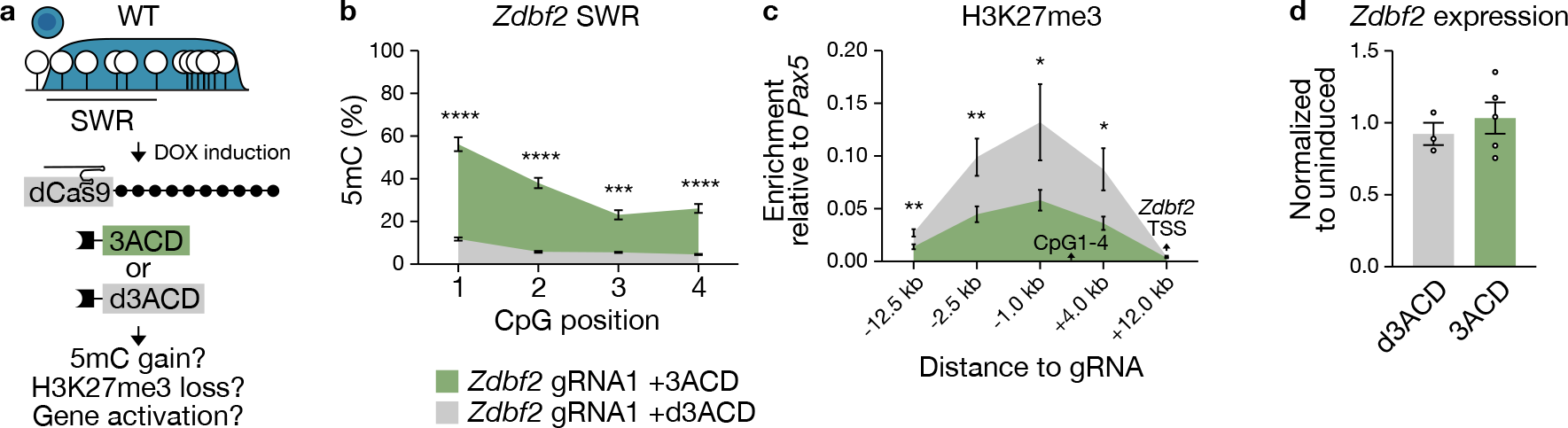
Site-directed 5mC deposition in naïve ESCs is sufficient to antagonize H3K27me3 at the *Zdbf2* SWR. **a.** Schema of induced targeted 5mC at candidate hypomethylated (white lollipops), H3K27me3 enriched (blue) SWRs. A construct composed of catalytically dead Cas9 (dCas9) and 10 GCN4 peptides (black circles) recruits multiple copies of the mouse DNMT3A catalytic domain (3ACD, green box) via single chain variable fragment antibodies (black squares). Catalytically dead DNMT3A (d3ACD, grey box) is used as a control. The construct is induced in naïve, globally hypomethylated ESCs and targeted to the *Zdbf2* SWR by gRNAs. 5mC, H3K27me3 and expression levels were assessed seven days post-induction. Not shown: GFP fused to 3ACD/d3ACD. **b.** Bisulphite-pyrosequencing (BS-pyro) showing 5mC levels at individual CpGs within the candidate *Zdbf2* SWR in cells expressing 3ACD (green, 9 replicates) and d3ACD (grey, 4 replicates). **c.** H3K27me3 CUT&RUN-qPCR levels normalized to the positive control *Pax5* CGI promoter (ΔCt method). Data represent 8 and 5 replicates for 3ACD and d3ACD induced samples, respectively. **d.** Bar plot showing *Zdbf2* expression levels measured by RT-qPCR. Levels were normalized to the average Ct of two housekeeping genes (*Rplp0* and *Rrm2*) and subsequently normalized to uninduced sample expression levels (∆∆Ct method). Data represent 5 and 3 replicates for 3ACD and d3ACD, respectively, denoted by dots. **b-d**. Data are shown as mean ± standard error. p-values were calculated by two-tailed unpaired t-test assuming equal variance: *p<0.05, **p<0.01, **p<0.001, ****p<0.0001.

To assess the efficiency of our epigenome editing system, we targeted the *Zdbf2* SWR as a proof-of-principle. We individually tested the extent of 5mC induction using four individual sgRNAs (*Zdbf2* gRNA1-4) followed by targeted deep methylation sequencing, with one guide (gRNA1) showing a 11.8% increase in 5mC levels spanning ∼2kb adjacent to the sgRNA target site (Fig. S9). Subsequently, dCas9-SunTag/3ACD induction experiments were performed in hypomethylated ESCs cultured in titrated 2i with the hope that this modified media composition would help stabilize the maintenance of newly deposited 5mC^46,70,71^. Using bisulfite treatment followed by pyrosequencing (BS-pyro), we obtained robust and highly reproducible 5mC deposition at the *Zdbf2* SWR (22-29%) compared to cells expressing d3ACD or uninduced controls (Fig. 4b & Fig. S10a).

While the extent of 5mC induction was modest, we nevertheless assessed the effect on H3K27me3 enrichment, by Cleavage Under Targets & Release Under Nuclease (CUT&RUN)^72^, followed by qPCR (Fig. 4c & Fig. S10b-c). Notably, H3K27me3 levels significantly decreased across a ∼16 kb region adjacent to the sgRNA target site in ESCs expressing dCas9-SunTag/3ACD compared to d3ACD. Despite loss of H3K27me3, *Zdbf2* expression levels were not significantly altered in epigenome edited cells (Fig. 4d). This may be due to incomplete depletion of H3K27me3 and/or lack of activating transcription factors in the naïve state, where *Zdbf2* is not typically expressed. Nevertheless, we were very encouraged by our ability to demonstrate direct antagonism of 5mC against PRC2 using a targeted approach.

We next selected four uncharacterized SWR regions for DNA methylation editing that looked promising from their genomic profiles: *Celsr2, Pdyn, Pga5* and *Krt80* (Fig 3d & Fig. S7d). Unfortunately, applying the same strategy did not yield a significant increase in 5mC levels (Fig. S10d). In total, 1/20 sgRNAs successfully induced 5mC gain at a target SWR, which could be due to the cell type, media conditions and/or the chromatin state of the loci we were targeting (see Discussion). Given these technical challenges, we took a complementary approach to DNA methylation editing.

### Site-specific DNA erasure during differentiation prevents H3K27me3 restriction, aberrantly maintaining PRC2-mediated repression

For our second approach, our goal was to perform locus-specific 5mC erasure in EpiLCs. We initially used the same inducible system described above, replacing the 3ACD with the TEN-ELEVEN-TRANSLOCATION 1 catalytic domain (TET1CD), which progressively oxidizes 5mC leading to 5mC loss, thus counteracting *de novo* 5mC deposition at target sites (Fig. S11a-b)^73^. However, once again we were met with a technical challenge, as our induction strategy could not overcome transgene silencing in differentiated cells (Fig. S11c). To overcome this limitation, we integrated a constitutively expressed dCas9-SunTag/TET1CD construct using piggyBac transposition (Fig. 5a, S11d)^74^. In this manner, robust expression of the construct was observed in the majority of cells after 4 days of differentiation (Fig. S11c); GFP positive cells were then enriched by fluorescent activated cell sorting.

**Figure 5.**
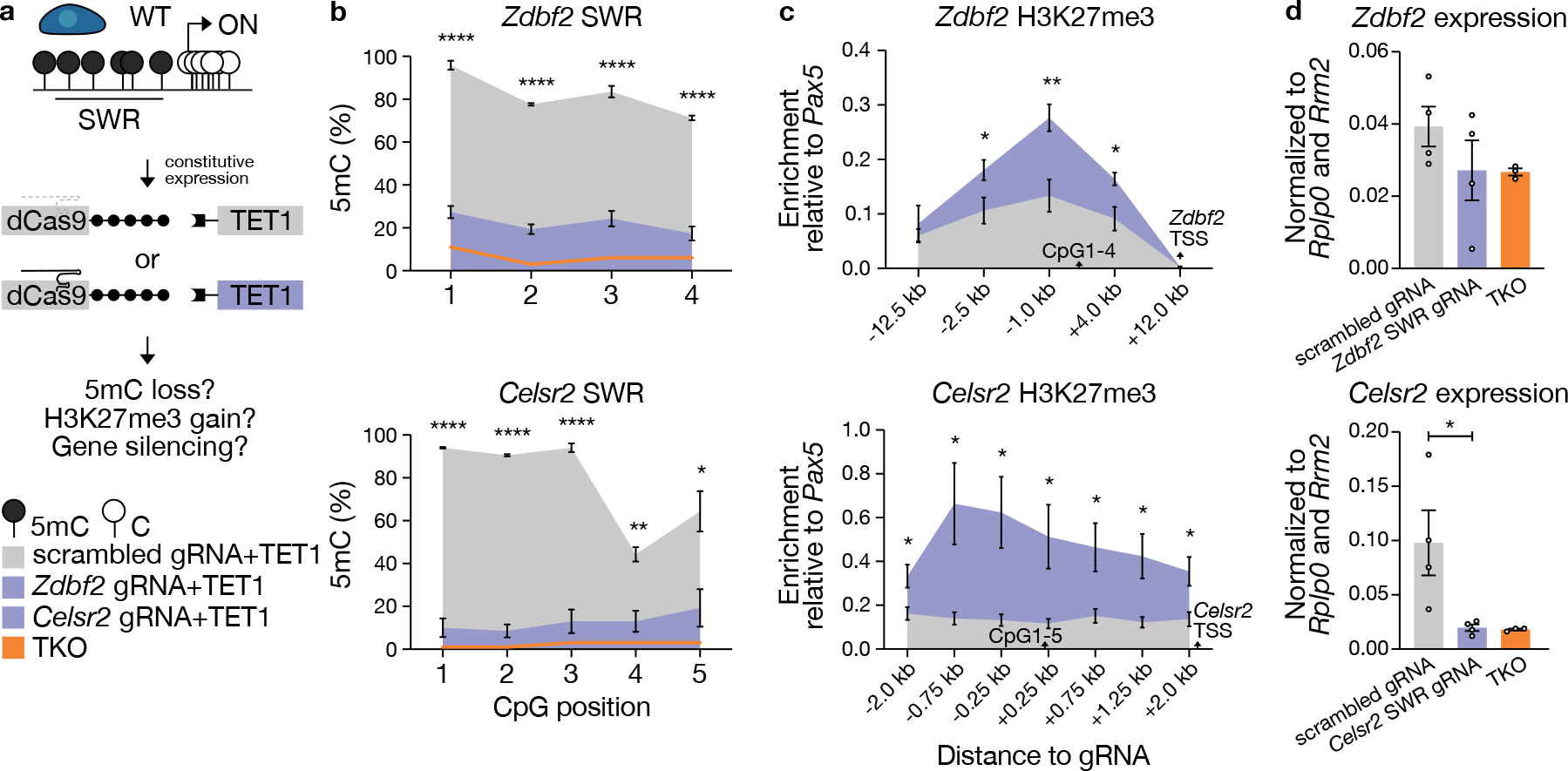
Targeted demethylation of candidate SWRs in differentiating ESCs leads to H3K27me3 maintenance and failure to activate *Zdbf2* and *Celsr2*. **a.** Schema of site-directed 5mC erasure at candidate SWRs. Demethylation is carried out by the catalytic domain of the human TEN-ELEVEN-TRANSLOCASE 1 (TET1CD) (purple). Site-specific demethylation is achieved by constitutive expression of dCas9-SunTag/scFv-TET1CD and gRNAs targeted to candidate SWRs. Cells constitutively expressing scrambled gRNAs were used as a control (grey). 5mC, H3K27me3 and expression levels were assessed in EpiLCs four days post-AF treatment. **b.** BS-pyro showing 5mC levels at individual CpGs within the *Zdbf2* (top) and *Celsr2* (bottom) candidate SWRs. Purple: targeted gRNAs, grey: scrambled gRNA. Data represent 4 replicates for each line. 5mC levels in TKO EpiLCs are included (orange, 1 replicate). **c.** H3K27me3 CUT&RUN-qPCR levels normalized to the positive control *Pax5* CGI promoter (ΔCt method) for the *Zdbf2* (top) and *Celsr2* (bottom) SWRs. Data represent 6 replicates for each line. **d.** Bar plot showing *Zdbf2* (top) and *Celsr2* (bottom) expression levels by RT-qPCR. Levels were normalized to the average Ct of two housekeeping genes (*Rplp0* and *Rrm2*) (ΔCt method). Expression levels in TKO EpiLCs are included (orange). 3-4 replicates were conducted for each sample, illustrated by dots. **b-d** data are shown as mean ± standard error. p-values were calculated by two-tailed unpaired t-test assuming equal variance: *p<0.05, **p<0.01, **p<0.001, ****p<0.0001.

Employing the optimized approach, we obtained remarkably robust and highly significant decrease in 5mC levels for all targets in EpiLCs: 60%, 65%, 38%, 61% and 75% decrease in 5mC relative to control cells for *Zdbf2, Celsr2, Pdyn, Pga5* and *Krt80*, respectively (Fig. 5b and Fig. S12a). Having achieved targeted DNA demethylation, we next set out to determine the impact on H3K27me3 using CUT&RUN-qPCR. For both *Zdbf2* and *Celsr2*, we observed a significant enrichment of H3K27me3 in epigenome edited cells compared to scrambled controls (Fig. 5c). The candidate SWR near *Krt80* showed a modest decrease in H3K27me3 levels in edited cells, and those near *Pdyn* and *Pga5* show depletion of H3K27me3 regardless of 5mC editing, suggesting that H3K27me3 depletion is uncoupled from 5mC deposition at the respective loci (Fig. S12b-c).

Remarkably, *Zdbf2* and *Celsr2*, the two genes that showed a clear maintenance of H3K27me3 levels when 5mC was actively removed, were downregulated in epigenome-edited cells to levels similar to those observed in TKO EpiLCs (Fig. 5d). In contrast, targeted 5mC demethylation of *Pdyn, Pga5 and Krt80* did not lead to relative repression at two different time points (Fig. S13a-b). Altogether, these results demonstrate that 5mC deposition is sufficient to antagonize PRC2-mediated repression at *Zdbf2* and *Celsr2*. In addition to validating our epigenome editing approach at the proof-of-principle locus, *Zdbf2*, we were able to identify *Celsr2* as a novel gene that is activated in a DNA methylation-dependent manner. Importantly, using a targeted approach also allowed us to distinguish *bona fide* SWRs from loci that exhibited SWR-like behavior in TKO EpiLCs that was likely simply coincidental. In sum, our TET editing strategy emerged as a promising means to demonstrate antagonism between two chromatin pathways typically associated with gene silencing, that in certain cases can lead to a change in gene-regulatory output. It also underscores the power of epigenome editing to discern locus-specific chromatin regulation from confounding effects that frequently occur in constitutive knockouts.

## Discussion

In this study we assessed the impact of 5mC deposition on shaping the H3K27me3 landscape in a cell-based model of embryonic progression, leveraging our prior finding that naïve mouse ESCs can differentiate to EpiLCs in the absence of DNA methylation machinery. Given the well-established antagonism between 5mC and PRC2, we hypothesized that in the absence of DNA methylation, the broad H3K27me3 domains characteristic of hypomethylated naïve ESCs would persist. We were somewhat surprised to discover a fairly typical, restricted H3K27me3 landscape in TKO EpiLCs. We went on to show that this was largely due to aberrant expression of the PRC2 antagonist *Ezhip*. Nevertheless, a substantial number of regions that switched from H3K27me3 enrichment to 5mC in WT EpiLCs, which we refer to as SWRs, maintained the PRC2 mark in TKO EpiLCs. Our studies of the *Zdbf2* locus demonstrated that embryonic DNA methylation deposition is required for H3K27me3 depletion and permissive expression^33^. Combining genome-wide analyses with precision epigenome editing, we discovered that *Celsr2* also requires DNA methylation establishment for Polycomb eviction and gene activation, suggesting that this is a wider form of gene regulation in early mammalian development.

Naïve ESCs exhibit elevated PRC2 protein abundance and pervasive H3K27me3 distribution throughout the genome, reinforced by PRC2-mediated repression of *Ezhip*. Indeed, the CGI promoter of *Ezhip* is within an ESR that normally maintains gene repression during normal differentiation. Consistently, *in vivo* post-implantation epiblasts do not express *Ezhip*. However, our differentiation experiments in QKO cells did not suggest an obvious impairment in transitioning from the naïve state—aside from a more “naïve-like” H3K27me3 profile—raising questions about the importance of H3K27me3 restriction as the cells prepare for lineage specification. It should be noted that there were perhaps more subtle differentiation or expression defects that we did not detect.

Primordial germ cells (PGCs), a cell type with low levels of DNA methylation, serve as an informative counterpoint to naïve ESCs. Mouse PGCs exhibit even lower 5mC levels than ICM cells^30,75^, and while not pluripotent, they share features with ESCs including expression of core pluripotency transcription factors^76^. A key distinction between the cell types is that unlike in ESCs, *Ezhip* is expressed in PGCs. Moreover, mice mutant for *Ezhip* exhibit increased H3K27me3 in both the male and female germline, which is associated with decreased fertility^60^. These observations lead us to propose that the CGI promoter of *Ezhip* acts as a global hypomethylation sensor that can help restrain potentially problematic H3K27me3 spreading. Curiously, this adaptation is not exploited in naïve ESCs. Does this imply that broad H3K27me3 domains are an important feature of the naïve state? Pertinent experiments for future studies could be to observe if ectopic *Ezhip* expression influences naïve ESC behavior by restricting the H3K27me3 landscape. Of course, this would need to be reconciled with the minimal impact on the naïve state in complete absence of PRC2 and H3K27me3^37,77,78^.

Even with derepressed *Ezhip* in TKO EpiLCs, we still observed a substantial number of H3K27me3-enriched regions in the genome that appear to be sensitive to deposition of DNA methylation in WT: SWRs. A probable important feature of SWRs is their intermediate CpG content. They seemingly occupy middle ground wherein they have exceeded a CpG density threshold that makes them attractive for PRC2 activity, but have not exceeded a threshold that would subject them to CGI regulation. With some notable exceptions, promoter CGIs exhibit several layers of protection—such as transcription factor binding^79,80^, H3K4me3^81–84^, and TET protein enrichment^85,86^—that keep them DNA methylation-free. And evidently at SWRs, the presence of H3K27me3 alone is not sufficient to deter *de novo* DNA methylation. What then could explain the sensitivity of PRC2 to 5mC? A potential interface could be the Polycomb-like (PCL) proteins— PHF1, MTF2 and PHF19—which are partially redundant components of the PRC2.1 subcomplex, and have been shown to be sensitive to DNA methylation^23,24^. It is important to note that PCL proteins are more strongly associated with CGI binding; however, profiling experiments were performed in highly methylated (*i*.*e*., serum-grown, ∼70% global 5mC) mouse ESCs, which would potentially preclude PCL protein presence at SWRs^23,24,87,88^. Thus, formal demonstration of PCL protein enrichment, or lack thereof, at SWRs in hypomethylated ESCs awaits.

While the primary focus of this study has been on PRC2-deposited H3K27me3, as previously mentioned, PRC1 and PRC2 activities are strongly interlinked^49^. The PRC1-associated RING1A and 1B proteins catalyze H2AK119ub, and published data in mouse ESCs indicates the vast majority of SWRs are enriched for this mark (Fig. S14a)^89^. Moreover, after two days of EpiLC differentiation, there already appears to be a shift towards H2AK119ub-marked SWRs in conditional *Dnmt* TKOs relative to WT (Fig. S14b-c). *De novo* DNA methylation is not complete at this time point, which would explain the relatively mild effect compared to our day seven H3K27me3 data (Fig. 1c). Canonically, H3K27me3 deposition serves as a platform for PRC1 binding; however, in mammals a number of non-canonical PRC1 (ncPRC1) complexes exist, which can bind to chromatin independently of PRC2 activity^90^. Notably, ncPRC1.1 and ncPRC1.6 contain 5mC-sensitive DNA binding factors KDM2B and MAX/MGA, respectively^25,91^. Publicly available ESC data indicate that PCGF2 (cPRC1.2 component) and PCGF6 (ncPRC1.6 component) exhibit the most overlap with SWRs (Fig. S14d)^66^. Most intriguingly, ncPRC1.6 is required for H3K27me3-to-5mC switching at a number of germline-specific gene promoters, although in these cases, the switch leads to maintained silencing^89,92,93^. It will be worthwhile to pursue a potential link between ncPRC1.6 and SWRs, and the possibility that ncPRC1.6 could facilitate gene activation via the same epigenetic switch mechanism that occurs at germline genes.

The development of site-directed epigenome editing has enabled testing the functional relevance of chromatin marks on local gene expression while bypassing traditional loss-of-function genetics of epigenetic regulators that can have far-reaching and confounding effects^94^. Initially, we attempted to establish 5mC using a dCas9-SunTag/3ACD system similar to ones already successfully implemented by other groups^95,96^. However, we had substantial difficulty establishing DNA methylation at SWRs, despite a great deal of effort and troubleshooting beyond using different combinations of sgRNAs; this included testing of alternative DNA methyltransferase domains, altering media composition, and using PRC2 inhibitors (data not shown). The resistance to targeted 5mC at these loci could be explained by a combination of factors acting against, or at least not favoring, the maintenance of the mark at the candidate regions in our cell system. Naïve ESCs are cultured in conditions with impaired DNA methylation maintenance machinery and high TET protein activity^97^. Even though DNA methylation maintenance machinery should be more stable in titrated 2i compared to 2i+vitC^71^, this still may not be sufficient for maintaining 5mC at our regions of interest. To wit, the nature of the sequences we targeted most likely do not contain features that bolster DNA methylation maintenance in the naïve state, such as those found at imprint control regions or endogenous retroviruses^98^. For example, the dCas9-SunTag/3ACD system successfully targeted 5mC at the *Dlk1-Dio3* imprint control region^99^ in mouse ESCs. While the establishment and maintenance of 5mC is enhanced with simultaneous targeting of a KRAB repressive domain^100^, this strategy was not appropriate in our system, as we were aiming to assess if DNA methylation led to gene activation. Finally, it should be noted that at least one guide RNA was able to reliably establish DNA methylation at the *Zdbf2* SWR, hence it is not an impossible task given a suitable guide and target sequence.

Conversely, the dCas9-SunTag/TET1CD system worked robustly and reliably at all candidate SWRs, enabling us to test if there was a regulatory role for DNA methylation at the underlying genes. Three of the candidates (*Pdyn, Pga5* and *Krt80*) did not exhibit relative repression in the absence of DNA methylation, and H3K27me3 levels were depleted in WT differentiated cells in the presence or absence of 5mC (Fig. S12c). This suggests that DNA methylation does not play a direct role at these regions, and underscores the hazard of making conclusions about 5mC-mediated regulation at specific loci when utilizing a constitutive knockout. However, we were encouraged by our demonstration that *Celsr2* genuinely requires DNA methylation to activate, validating our approach. It is worth noting that we only analyzed 5/68 candidate genes using epigenome editing, so there are certainly more opportunities to discover similar examples of this mode of gene control. There are also two important points to stress: we only focused on genes where we could observe differential gene expression in WT and TKO EpiLCs, but there could be a much larger set of genes that we omitted due to their activation at later time points in development. This is a limitation of using TKO cells, which do not exhibit the same differentiation potential as WT. Further, for simplicity we focused on SWRs that overlapped gene promoters, but a similar mode of regulation could occur at intergenic *cis* regulatory elements, such as enhancers. Indeed, DNA methylation has been suggested to be important to activate enhancers in some contexts^101^. Both points can be fodder for future studies.

The *Celsr2* gene encodes an atypical cadherin protein that is widely expressed in the developing and adult mouse and human central nervous systems^102–106^. Knockout studies indicate pleiotropic phenotypes, including hydrocephaly^107^ and more recently, a decrease in cortical plasticity and motor learning in mice^104^ and an increase of motor neuron regeneration after injury in mouse and human explants^108^. To our knowledge, there is no early embryonic phenotype associated with *Celsr2* KOs. This creates the distinct possibility, that like *Zdbf2*, it is the embryonic DNA methylation programming that permits *Celsr2* activation in later developmental timepoints. Consistently, publicly available data^109^ indicates that *Celsr2* maintains its high 5mC levels in tissues where it is highly expressed (Fig. S14e). With emerging *in vivo* TET-mediated epigenome editing techniques^74,110^, it will be exciting to interrogate the relevance of 5mC-mediated Polycomb antagonism in a physiologically relevant system.

The combined power of genome-wide analyses with single-locus epigenome modulation opens doors to discover and characterize atypical modes of chromatin regulation, not only in the context of development as described here, but also in disease. Epigenome misregulation is characteristic of virtually all cancers, and SWR regions are prevalent^12,111,112^—although it is difficult to discern cause from consequence. With epigenome editing, it is now possible to test whether SWRs are associated with a change in gene expression, and possibly disease progression.

## Materials & Methods

### Mouse embryonic stem cell culture

E14Tg2a (E14) mouse ESCs was the parental line used for all experiments in this study, as well as serving as the background for all transgenic lines. The E14 *DnmtTKO* line was previously generated in-house ^47^.

For the cells grown in serum culture conditions we used Glasgow medium (Gibco) supplemented with 15% Fetal bovine Serum (FBS), 0.1 mM MEM non-essential amino acid, 1mM Sodium Pyruvate, 2mM L-Glutamine, Penicillin/Streptomycin, 0.1 mM β-mercaptoethanol and 1000 U/mL Leukemia Inhibitory Factor (LIF). To pass, the cells were washed with 1X PBS, then trypsin was added to detach and disaggregate the cells for 5 minutes at 37°C. The desired number of cells were then transferred to the new flask.

For the 2i+vitC culture conditions (naïve ESCs) we used N2B27 medium (50% neurobasal medium, 50% DMEM) supplemented with N2 (Gibco), B27 (Gibco), 2 mM L-Glutamine, 0,1 mM β-mercaptoethanol, Penicillin/Streptomycin, LIF and 2i (3 μM Gsk3 inhibitor CT-99021, 1 μM MEK inhibitor PD0325901) and Vitamin C (Sigma) at a final concentration of 100 μg/mL. Titrated 2i (t2i) medium was prepared as 2i medium with two modifications: the MEK inhibitor PD0325901 was used at 0.2 μM and no Vitamin C was supplemented. To pass the cells, the media was removed, then Accutase (Gibco) was added to detach and disaggregate the cells and incubated for 5 minutes at room temperature. The desired number of cells were then transferred to the new plate. The ESCs in all three conditions were grown on 0.1% gelatin-coated flask in an incubator at 37°C and 5% CO_2_.

To induce EpiLC differentiation, cells were gently washed with PBS, dissociated, and replated at a density of 2 × 10^5^ cells/cm^2^ on Fibronectin (10 μg/mL, Sigma)-coated plates in N2B27 medium supplemented with 20 ng/mL Activin A (R&D) and 12 ng/mL FGF2 (R&D) (AF treatment). EpiLCs were passed with Accutase on day 2 or 3 of differentiation depending on the end time-point of differentiation: day 4 or 7, respectively.

Mycoplasma contamination checks were performed using an ultrasensitive qPCR assay (Eurofins).

### Plasmid Generation

The Dox-inducible dCas9-SunTag is a modified version of the pHRdSV40-dCas9-10xGCN4_v4-P2A-BFP vector (Addgene #60903). An N-terminal DD domain and the construct minus the P2A-BFP were PCR amplified and Gibson cloned into a version of the pEN111 vector (a gift from Elphege Nora) containing a Neomycin resistance gene. The Dox-inducible scFv-GFP-3ACD, d3ACD, or TET1CD was generated as follows: pHRdSV40-scFv-GCN4-sfGFP-VP64-GB1-NLS (Addgene #60904) was digested with *Rsr*II and *Spe*I restriction enzymes to remove the VP64 sequence, as previously performed^95^. Then either the 3ACD, a d3A CD containing a Cysteine-to-Alanine mutation in the catalytic loop, or the TET1CD was inserted by Gibson cloning. Subsequently, an N-terminal DD domain and the construct were PCR amplified and inserted into the original pEN111 vector containing puromycin resistance gene by Gibson cloning.

All CRISPR/Cas9 guide sequences were cloned into a piggyBac donor plasmid containing a U6 promoter and enhanced gRNA scaffold sequence and downstream hygromycin resistance gene^113^. Single guides were cloned by *Bbs*I digest and ligation of double stranded guide sequences with compatible overhangs. Dual guides were cloned by amplifying a PCR product from the pLKO.1-blast-U6-sgRNA-BfuA1-stuffer plasmid^6^ containing one gRNA sequence, the enhanced gRNA scaffold, a modified mouse U6 promoter^114^ and a second gRNA sequence, and inserting it into the *Bbs*I digested piggyBac donor vector by Gibson cloning. All guide sequences were designed using the CRISPOR web tool (crispor.tefor.net) and are listed in Supplemental Table 2. Epigenome editing gRNA target sites were determined based on previously published data on RING1B and SUZ12 occupancy^34^.

### Ezhip deletion strategy

The *Ezhip* deletion was generated by transfecting SpCas9 and two CRISPR single guide RNAs specific to target sequences flanking the entire *Ezhip* coding sequence. Guide sequences were designed using the online CRISPOR online program (crispor.tefor.net) and cloned into the pX459 plasmid harboring the *Cas9* gene. All guide sequences are listed in Supplemental table 2. Transfections were performed in *Dnmt* TKO ESCs cultured in serum using Lipofectamine 2000 (Life Technologies) according to the manufacturers instructions. Following a two-day treatment of puromycin to select for transfectants, 96 individual clones were picked and screened by PCR for deletion. Mutated alleles were confirmed by Sanger sequencing of cloned PCR amplicons, and two *Ezhip* KO subclones were kept for analysis. All primer sequences are listed in Supplemental table 2.

### Generation of stable epigenome editing lines

The Dox-inducible epigenome editing lines were generated as follows: The pEN111-DD-dCas9-SunTag vector (described above) was co-transfected into serum-grown ESCs along with a pX330-EN479 (Addgene #86234), which contains a Cas9 nickase and a guide targeting the *Rosa26* locus, using Lipofectamine 2000 (Life Technologies) according to the manufacturer’s instructions. After one day, Genetecin (Gibco) was added to media at a final concentration of 250 μg/mL. Approximately one week later, 96 individual Neomycin resistant clones were picked and screened for hemizygous insertion at *Rosa26* by PCR genotyping. The unmodified alleles of positive clones were analyzed by Sanger sequencing to ensure that there was no CRISPR/Cas9-induced mutations at the guide target site. Finally, inducible expression was confirmed by western blot. The scFv-GFP-3ACD/d3ACD/TET1CD constructs were transfected using the same method into a dCas9-SunTag/+ line, using puromycin (Gibco) at a final concentration of 1μg/mL. Induction was confirmed by western blot and fluorescent microscopy.

All piggyBac donor plasmids were co-transfected with pBroad3_PBase_IRES_RFP (a gift from Luca Giorgetti), which contains the piggyBac transposase, using Lipofectamine 2000. For guide constructs, stable integration was confirmed by Hygromycin (Invitrogen) selection at a final concentration of 200 μg/mL. Stable integration of the constitutive dCas9-SunTag/TET1CD construct was confirmed by fluorescence-activated cell sorting (FACS) of GFP positive cells with a Becton Dickinson FACSAria™ Fusion after 10 days.

### Epigenome editing

For 3ACD and d3ACD experiments, cells were grown in 2i+vitC medium for at least seven days before switching to t2i conditions. To induce the expression of the epigenome editing machinery, Doxycyclin (Sigma) and Shield-1 (Aobious) were added to the medium at a final concentration of 0.5 μg/mL and 100nM, respectively, for seven days prior to collection of the cells.

For inducible TET1CD experiments, cells were grown in 2i+vitC medium for at least seven days before starting the differentiation to EpiLC. The induction was performed as above, but only for four days and in parallel with EpiLC differentiation.

For constitutive TET1CD experiments, after the line was generated and confirmed, serum-grown ESCs were converted to 2i+vitC for 10 days, and then differentiated to EpiLCs. For both the constitutive and the inducible TET1CD lines, cells were FACS sorted for GFP expression at day 4 or 7 of EpiLC differentiation.

### Targeted Deep Methylation Sequencing

DNA was converted using NEBNext Enzymatic Methyl-seq conversion module (New England Biolabs) according to the manufacturer’s instructions. We used this technique because the damage caused to DNA is minor compared to bisulfite treatment and DNA residues are changed in the same manner, thus minimally affecting downstream analyses. Briefly, the genomic DNA was first sheared by sonication with Bioruptor. Then, 420ng of shared DNA were used for conversion. The 5-methylcytosines and the 5-hydroxymethylcytosines were oxidized by the TET2 protein included in the kit. The oxidized DNA was further deaminated and purified.

We proceeded to generate material following the protocol described in Leitão et al.^115^. The converted DNA was amplified with two rounds of PCRs. The first round includes forward and reverse universal tags. We designed primers for 6 amplicons of ∼300bp covering a ∼2kb region encompassing the potential polycomb nucleation site upstream *Zdbf2* transcription start site.

After amplification the DNA was quantified by Qubit and the 6 amplicons were pooled in an equimolar manner for each sample. Each pool was then purified and the pools were amplified in a second round of PCR to include Illumina-compatible sequences. The 2nd PCRs were quantified with Qubit and their quality and size were checked with a Bioanalyzer (Agilent). The 20 PCRs corresponding to the 20 different samples were pooled in an equimolar manner and purified twice with the SPRI based size selection following the kit instructions for selection of the fragment between 400 to 1000bp. This purified pool was then quantified and run in the Bioanalyzer. Finally, the sample was prepared at a concentration of 4nM containing 10% of PhiX DNA and 10% of another library provided by colleagues do add sequence complexity to the pool in order to facilitate the discrimination of clusters. Paired-end, 250 bp sequencing was performed on an Illumina miSeq machine. All primers are listed in (Supplemental Table 2).

### Bisulfite-Pyrosequencing

Genomic DNA was isolated from cells using the NucleoSpin Tissue kit (Macherey-Nagel). 500 ng of genomic DNA was subject to bisulfite convertion using the EZ DNA Methylation-Gold kit (Zymo). Target genomic regions were amplified by PCR using 1-2 μL of converted DNA with specific primers pairs recognizing BS-converted DNA, one of which being biotin-conjugated (Supplemental Table 2), using Immolase (Ozyme) or GoTaq® G2 (Promega) polymerases. Pyrosequencing assays were designed using the PyroMark Q24 Advanced 3.0.1 software and the sequencing reaction was performed with PyroMark Q24 Advanced reagents (Qiagen, 970922) according to manufacturer’s instructions. Briefly, 20 μL of the PCR reaction was mixed with streptavidin beads (Cytiva, 17511301) and binding buffer, by shaking for 10 min at RT. Samples were then denaturated with denaturation buffer using a PyroMark workstation (Qiagen) and released in PyroMark Q24 plate (Qiagen) pre-loaded with 0.375 μM of sequencing primer. The single-strand PCR template was annealed to the sequencing primer by heating at 80ºC for 5 min. The plate was then processed in the PyroMark Q24 pyrosequencer (Qiagen). Results were analyzed with PyroMark Q24 Advanced 3.0.1 software. Graphical representation and statistical analysis was performed with GraphPad Prism v9.3.1.

### CUT&RUN-qPCR

To measure H3K27me3 enrichment after epigenome editing, we used the original CUT&RUN l protocol^72^ with minor modifications. Briefly, 2.5-5 x 10^5^ cells per sample were pelleted and washed twice with wash buffer (20 mM HEPES pH 7.5, 150 mM NaCl, 0.5 mM spermidine containing protease inhibitor). The cells were then incubated for the binding with Concavalin A magnetic beads (Polysciences) by rotating for 10 min at RT. The antibody incubation was carried out overnight on a rotor at 4ºC in 100 μL antibody solution (wash buffer with 0.1% digitonin and 2 mM EDTA) containing 1 μL of αH3K27me3 Ab (Cell Signaling C36B11). Samples were then washed in dig-wash buffer (wash buffer with 0.1% digitonin), and incubated with pAG-Mnase on a rotor for 10 min followed by two more washes. The Mnase reaction was then activated by adding 2 mM CaCl_2_. After incubating at 0°C for 30 min the reaction was stopped with 1x final concentration of STOP solution (2x: 340 mM NaCl, 20 mM EDTA, 4 mM EGTA, 0.02% digitonin, 1:200 Rnase A). Chromatin was released by incubating samples at 37°C for 10 min. After centrifuging at full speed for 5 min, the supernatant was collected after incubation on magnet stand. Purified DNA was obtained by phenol/chloroform extraction and precipitated with 100% ethanol by centrifugation. The DNA pellet was washed in 70% ethanol, spun down and air-dried before being resuspended in 35 μl of 5 mM Tris-HCl pH 8.5. The pAG-Mnase plasmid was obtained from Addgene (# 123461), and the protein was purified by the Curiecoretech Recombinant Protein Platform.

CUT&RUN DNA fragments were then subjected to quantitative PCR to amplify selected target regions: 1 μL DNA was mixed with 5 μL of LightCycler 480 SYBR Green I Master Mix and 4 μL of forward and reverse primer mix (final primer concentration 200 nM). Primers specific for target and control regions (positive control: *Pax5* and negative control: *β-Actin*) were used. RT-qPCR was run on a LightCycler 480 II (Roche Applied Science) using 384 well plates (see RT-qPCR section for thermocycling details). Relative enrichment of H3K27me3 was calculated by comparing Ct values of target regions to the positive control locus, *Pax5* (ΔCt method). Graphical representation and statistical analysis were performed with GraphPad Prism v9.3.1. Primer sequences can be found in Supplemental Table 2.

### Protein Extraction & Western Blot

For protein extraction, cells were incubated with a BC250 lysis solution (25 mM Tris pH 7.9, 0.2 mM EDTA, 20% Glycerol, 0.25 M KCl) supplemented with protease inhibitors (Roche) and sonicated 3x5 seconds using a Bioruptor (Diagenode). The lysate was centrifuged at 13K g for ten minutes, and the supernatant was transferred to a new tube. Protein concentrations were quantified using Pierce BCA Protein Assay Kit (ThermoFisher) on an Infinite M200 plate reader (Tecan). For Western Blotting, 3 μg of the protein samples were prepared with 4X Laemmli loading buffer (Biorad) and denatured at 95°C for 5 minutes. The samples were then loaded in Mini-PROTEAN TGX precast gels (Biorad) and run at 100V for 15 minutes and 1 hour at 130V. Proteins were then transferred onto a PVDF membrane and rinsed with PBS-Tween (1%). The membrane was blocked in 5% milk in PBS-T for 30 minutes. Membranes were then cut and incubated with different primary antibodies diluted in 5% milk at 4°C overnight: αH3K27me3 (Cell Signaling C36B11) was used at 1:5000 dilution, and αLamin-B1 (abcam ab16048) at 1:2000. Blots were washed three times with PBST for 10 minutes at room temperature, and incubated for 1 hour with fluorescent-tagged secondary antibody diluted in PBST (Goat anti-Rabbit IgG StarBright™ Blue 700 or Goat anti-Mouse IgG DyLight™ 800, Bio-Rad). Finally, we performed 3 washes with PBST for 10 minutes before imaging the membrane on a ChemiDoc system (BioRad). The original scans of the blots are provided in Supplemental Table 3.

### LUMA

500 ng genomic DNA was digested with MspI+EcoRI or HpaII+EcoRI (New England BioLabs). HpaII is a methylation-sensitive restriction enzyme, and MspI is its methylation-insensitive isoschizomer. EcoRI was included for internal normalization. The extent of the enzymatic digestions was quantified by pyrosequencing (PyroMark Q24), and global CpG methylation levels were then calculated from the HpaII/MspI normalized peak height ratio.

### RT-qPCR

Total RNA was extracted from cell pellets using NucleoSpin RNA kit (Macherey-Nagel). First-Strand cDNA synthesis was performed using the SuperScript III Reverse Transcriptase kit (Invitrogen). 500-1000 ng of total RNA per reaction were mixed with 1 μL 50 ng/μL of random primers, 1 μL of 10 mM dNTPs and sterile H_2_0 up to 13 μL. The obtained cDNA was diluted 1:5 and for each RT-qPCR reaction 1 μL cDNA was mixed with 5 μL of LightCycler 480 SYBR Green I Master Mix and 4 μL of forward and reverse primer mix (final primer concentration 200 nM). RT-qPCR was run on a LightCycler 480 II (Roche Applied Science) using 384 well plates. The samples first followed an initial incubation at 95°C for 10 min, and then 45 cycles of denaturation at 95°C for 10 seconds, annealing at 61°C for 20 seconds and extension at 72°C for 20 seconds. Samples were amplified in triplicates with appropriate non-template controls. Relative gene expression was calculated using the 2ΔCt method and normalized to the geometric mean of the expression levels of the two housekeeping genes *Rrm2* and *Rplp0*. Graphical representation and statistical analysis were performed with GraphPad Prism v9.3.1. Primer sequences can be found in Supplemental Table 2.

### WGBS

From genomic DNA we provided, WGBS libraries were generated using the Accel-NGS Methyl-Seq DNA library Kit (Swift Biosciences) and sequenced on an Illumina Novaseq (paired-end, 150 nucleotide-long reads) by Novogene Co.

### CUT&TAG

We performed CUT&Tag according with the protocol described in ^45^, closely following the procedure described in “Bench top CUT&Tag V.2”^116^. 500,000 cells were used for each sample and duplicates were performed. The primaries antibodies, anti-H3K27me3 (Cell signal C36B11) 1:100, the anti-H3K36me3 (Invitrogene RM491) 1:100, were incubated overnight at 4ºC on a rotator. Samples were then incubated with a secondary anti-rabbit IgG (ABIN6923140) at room temperature on a rotator. After an incubation of 1h at room temperature with a pA-Tn5 adapter complex (Addgene #124601), the Tn5 enzyme tagmentation reaction was performed by adding a buffer containing MgCl_2_ to the samples and incubating for 1h at 37°C. To stop tagmentation and solubilize DNA fragments, we added 10 μL 0.5M EDTA, 3 μL 10% SDS and 2.5 μL 20 mg/mL Proteinase K to each sample and samples were incubated at 50°C for 1h. Purified DNA was obtained by phenol/chloroform extraction and precipitated with 100% ethanol and centrifugation. The DNA pellet was washed in 80% ethanol, spun down and air-dried before being resuspended in 25μl of 1mM Tris-HCl pH8.0. Nextera primers or indexed primers were used to perform PCR. Library quality was assessed using 4200 TapeStation system (Agilent) and paired-end, 150 bp sequencing on an Illumina Novaseq was performed by Novogene Co.

### High throughput sequencing read filtering and trimming

Prior to alignment, HTS library sequencing quality of was assessed using FastQC (v0.11.9)^117^. PCR duplicate reads were removed using BBMap clumpify (v38.18)^118^ and parameters:

*reorder dedupe=t k=19 passes=6 subs=(0*.*01 x readlength)*

Where the number of substitutions allowed is equal to the length of the read in nucleotides multiplied by the conservative sequencing error rate of Illumina machines. Finally, adapter and low quality sequences were trimmed using Trimmomatic (v0.39)^119^ and parameters:

*ILLUMINACLIP:adapters*.*fa:2:30:10*

*SLIDINGWINDOW:4:20 MINLEN:24*

Where adapters.fa corresponds to a fasta file containing commonly used Illumina sequencing adapters. H3K36me3 CUT&Tag data was processed with the additional parameter CROP:36.

### WGBS analysis

Filtered and trimmed (see above) HTS reads we’re aligned to the mm10 genome using Bismark (v0.23.1)^120^ and default parameters. Reads with mates that did not survive read trimming or that could not be aligned in paired-end mode were concatenated and realigned in single-end mode. PCR duplicate reads were removed using deduplicate_bismark and CpG methylation information was extracted using bismark_methylation_extractor. Average CpG methylation levels over regions of interest was calculated using BEDOPS (v2.4.40)^121^. For each dataset, 10kb bins, and CGIs with at least 10 CpGs covered by at least 5 reads were kept. H3K27me3 peaks with at least 5 CpGs covered by at least 5 reads were kept. Spaghetti and violin plots were made using VisR (v0.9.42)^122^.

### RNAseq analysis

Filtered and trimmed (see above) RNAseq reads we’re aligned to the mm10 genome using STAR (v2.7.9a)^123^ and default parameters. Transcript (UCSC RefGene) and gene level (NCBI RefSeq) expression values were calculated using VisR using uniquely aligned reads (MAPQ=255). *De novo* transcriptome assembly was performed for each sample using Stringtie (v2.1.7)^124^ with default parameters. *De novo* assemblies were subsequently combined and annotated using the UCSC RefGene reference file. The transcription start site of each isoform was extracted using a custom AWK script. Transcript level expression values (RPKM) were normalized for batch effects using pyCombat (v0.3.2)^125^ and input into sklearn PCA (v1.1.1)^126^ using pandas (v1.4.3)^127,128^ and numpy (v1.23.1)^129^ to generate a PCA plot which was visualized using matplotlib (v3.5.2)^130,131^ and seaborn (v0.11.2)^132^. Alignment files were converted to bigwig using Deeptools bamCoverage (v3.5.1)^133^ and parameters:

*--binSize 1 –smoothLength 0 –minMappingQuality 255 –normalizeUsing CPM –outFileFormat bigwig – blackListFileName ENCFF547MET*.*bed – ignoreForNormalization chrX chrM chrY*

Where ENCFF547MET.bed corresponds to the coordinates of blacklisted regions as defined by the Kundaje lab as part of the ENCODE consortium. Scatter plots and heatmaps of gene expression were generated in Python using matplotlib, numpy, pandas and seaborn. Correlograms were generated using Morpheus and spearman rank correlation of gene expression (RPKM) values. Linear modelling using Limma (v3.54.1)^134^ in R was employed to identify statistically significant differences in gene expression levels between samples. Voom normalization and a stringent threshold of log2FC>1 and adjusted p value (t-test) <0.05 was applied to define statistical enrichment.

### CUT&TAG data analysis

Filtered and trimmed (see above) CUT&TAG data was aligned to the mm10 reference using Bowtie2 (v2.4.5)^135^ and parameters:

*--local –very-sensitive –no-mixed –dovetail –no-discordant –phred33 -I 10 -X 700*

Replicate alignment files were merged prior to peak calling using the SEACR (v1.3)^136^ and parameters:

*0*.*01 non stringent*

All H3K27me3 peaks were merged using bedtools merge (2.30.0)^137^ and H3K27me3 enrichment levels (RPKM) were calculated using VisR from alignments with MAPQ ≥ 10. H3K27me3 domains were agnostically clustered into 6 groups using k-means (Morpheus, Broad Institute) of RPKM values. Statistical enrichment was performed as described above for RNA-seq using the same filtering criteria. Sunset plots and meta plots of H3K27me3 enrichment over groups were generated using Deeptools functions computeMatrix and plotHeatmap and plotGroup. The number of H3K27me3 peaks overlapping CGIs, the promoters of NCBI RefSeq genes (TSS +/-1kb) or bodies of genes was assessed using Bedtools intersect. Perkiness scores were calculated by averaging the enrichment level (RPKM) of the top 10% enriched peaks for each dataset, as previously described^27^. Total domain size was calculated by summing the lengths of all peaks for each dataset. Alignment files were converted to bigwig using Deeptools bamCoverage and the same parameters as for RNA-seq analysis (see above) with the exception of:

*--minMappingQuality 10*

Transcription factor binding motif analysis of SWRs and ESRs was performed using the MEME^138^ and Rsat^139,140^ suites.

### Measuring CpG density

A CpG density track was generated following previously described method^2^. Briefly, 100 bp overlapping 1000 bp bins were generated using Bedtools makewindows and the number of Cs, Gs, and CpGs were counted using Bedtools getfasta and faCount. CpG density was then calculated for each bin using the formula:

*CpG observed/expected ratio = #CpG / (#C + #G)*

The average CpG density ratio was calculated using bedmap, generating a bedGraph file of 100 bp windows, which was then converted into a genome browser compatible track using bedgraphtobigwig^141^.

### Data visualization

UCSC Genome Browser^142^ Track Hubs^143^ were used to display data over loci of interest.

## Declarations

### Availability of data and materials

Datasets generated in this study were uploaded to NCBI GEO under accession number GSE242201. See Supplemental Table 1 for the full list of data analyzed in this study. Scripts used to generate the figures presented are available under an GNU General Public License v3.0 on GitHub (https://github.com/julienrichardalbert/5mC-H3K27me3/releases/tag/v0.1).

### Authors’ contributions

**J**.**R**.**A**.: Conceptualization, Methodology, Formal analysis, Data Curation, Writing – Original Draft, Writing – Review & Editing, Visualization **T**.**U**.: Investigation, Writing – Original Draft, Writing – Review & Editing, Visualization **A**.**M-S**.: Investigation **A**.**L**.**B**.: Investigation **A**.**D**.: Investigation **M**.**S**.: Investigation **M**.**V**.**C**.**G**.: Conceptualization, Methodology, Investigation, Writing – Review & Editing, Supervision, Project administration, Funding acquisition

## Acknowledgements

Takuro Horii and Izuho Hatada for providing the constitutive dCas9-SunTag/TET1CD epigenome editing construct. Bérengère Guichard and Martin Möckel for purifying pA-Tn5 protein. Cara McQuillen and Hugo Fanlo Ucar for technical assistance. Déborah Bourc’his for her insights and support. Martin Leeb, Aaron Bogutz, Sanne Janssen, and Mattia Pitasi for useful discussions. Matthew Lorincz and Daniel Holoch for critical reading. Joël Marchand for installing and maintaining computational resources. Nicolas Valentin and the IJM FACS facility.

## Supplemental tables

Supplemental table 1: Datasets generated and mined in this study.

Supplemental table 2: Oligonucleotides used in this study.

Supplemental table 3: Source data.

---

**Figure S1.**
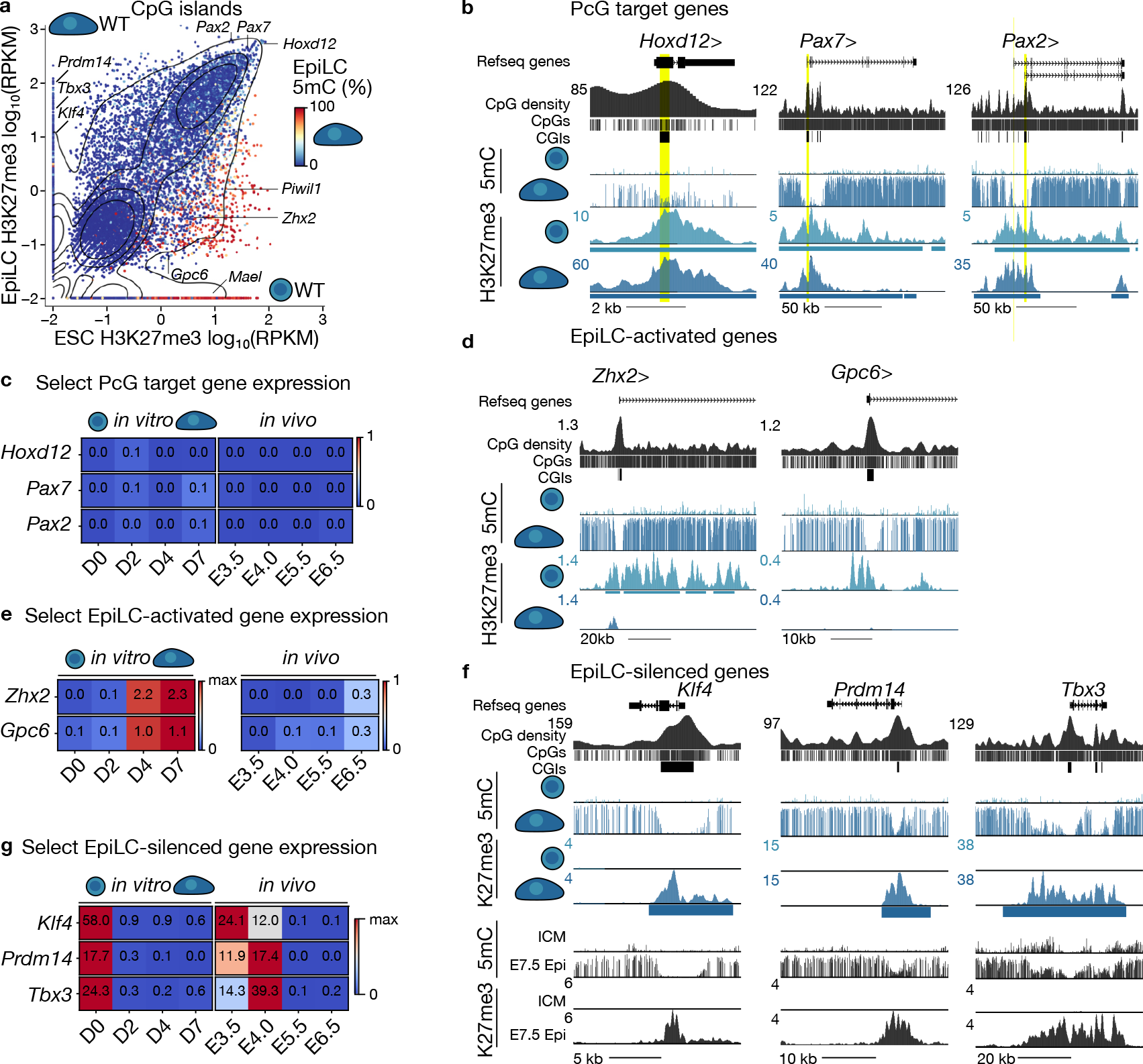
H3K27me3 dynamics over CpG islands during EpiLC differentiation. **a.** Scatterplot showing H3K27me3 levels over CpG islands (CGIs) in WT ESCs and EpiLCs (7 days post-Activin A/FGF2: AF treatment). Datapoints are coloured by average 5mC levels in WT EpiLCs. CGIs with ≥10 CpGs covered by ≥5 reads are shown (n=15,714/16,009). CGI promoters of specific genes are indicated, contours are drawn at iso-proportions of the density of datapoints. **b.** UCSC Genome Browser screenshots of Polycomb-group target genes. WGBS and H3K27me3 CUT&Tag data derived from WT ESCs and EpiLCs (7 days post-AF treatment) is shown. Refseq genes, CpG density, individual CpGs are included. Coordinates: *Hoxd12* chr2:74,672,861-74,678,432, *Pax7* chr4:139,712,945-139,857,645, *Pax2* chr19:44,712,108-44,849,206. **c.** Heatmap showing expression levels (RPKM) of Polycomb-group (PcG) target genes *Hoxd12, Pax7* and *Pax2 in vitro* (0, 2, 4 and 7 days post-AF treatment) and *in vivo* (inner cell mass cells of embryonic day (E) 3.5 and E4.0 blastocysts, and E5.5 and E6.5 epiblasts. **d.** UCSC Genome Browser screenshots of EpiLC-specific genes. Data as in **b**. Coordinates: *Zhx2* chr15:57,676,294-57,758,534, *Gpc6* chr14:116,904,924-116,947,162 **e.** Heatmap showing expression levels (RPKM) of EpiLC-specific genes *Zhx2* and *Gpc6*, as in **c**. For each *in vitro* and *in vivo* dataset, the scale bar for each gene is set independently, either to the maximum expression level or 1, whichever is higher. **f.** UCSC Genome Browser screenshots of naïve EpiLC-silenced genes. Data as in **b**. *in vivo* data derived from the inner cell mass (ICM) of E3.5 blastocysts and E7.5 epiblasts is shown. Coordinates: *Klf4* chr4:55,521,819-55,537,790, *Prdm14* chr1:13,104,889-13,135,730, *Tbx3* chr5:119,635,577-119,698,095. **g.** Heatmap showing expression levels (RPKM) of EpiLC-silenced genes *Klf4, Prdm14* and *Tbx3*, as in **c**.

**Figure S2.**
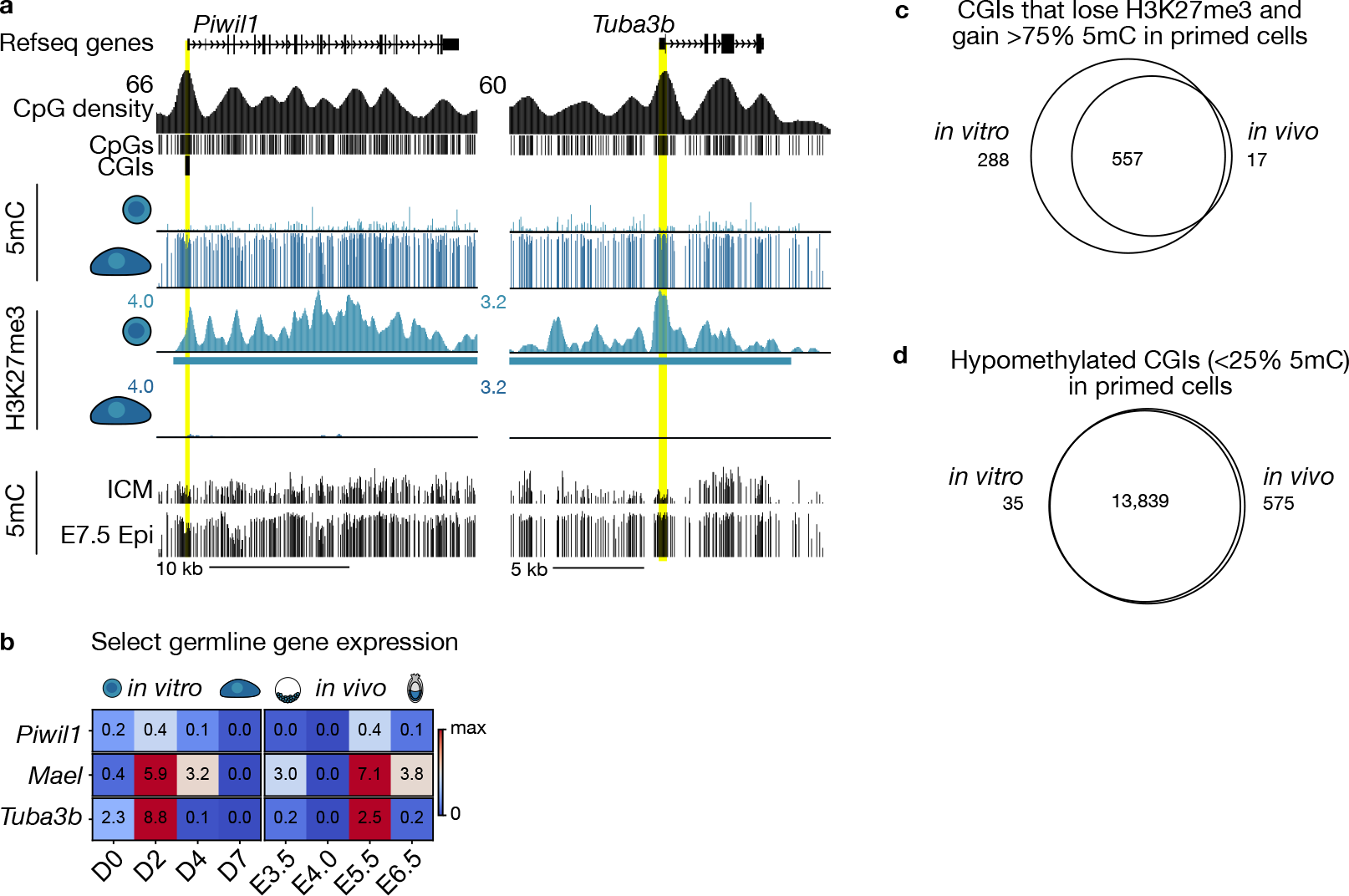
Germline genes remain repressed after a H3K27me3-to-5mC switch at their CpG-rich promoters during EpiLC differentiation. **a.** UCSC Genome Browser screenshots of germline-specific genes. WGBS and H3K27me3 CUT&Tag data derived from WT ESCs and EpiLCs (7 days post-AF treatment) is shown. Refseq genes, CpG density, individual CpGs are included. *In vivo* WGBS derived from the inner cell mass (ICM) of E3.5 blastocysts and E7.5 epiblasts are shown. Coordinates: *Piwil1* chr5:128,733,977-128,756,805, *Tuba3b* chr6:145,607,562-145,625,145. **b.** Heatmap showing expression levels (RPKM) of germline-specific genes *Piwil1* and *Tuba3b, in vitro* (0, 2, 4 and 7 days post-AF treatment) and *in vivo* (inner cell mass cells of E3.5 and E4.0 blastocysts, and E5.5 and E6.5 epiblasts). **c.** Venn diagram showing the overlap of CGIs that concomitantly gain 5mC (>75%) and lose H3K27me3 levels (linear modelling using Limma, FC>2, t-test adjusted p value<0.05) during exit from naïve pluripotency *in vitro* and *in vivo*. **d.** Venn diagram showing the overlap of hypomethylated CGIs (<25% 5mC) in EpiLCs *in vitro* (7 days post-AF treatment EpiLCs) and *in vivo* (E7.5 epiblasts).

**Figure S3.**
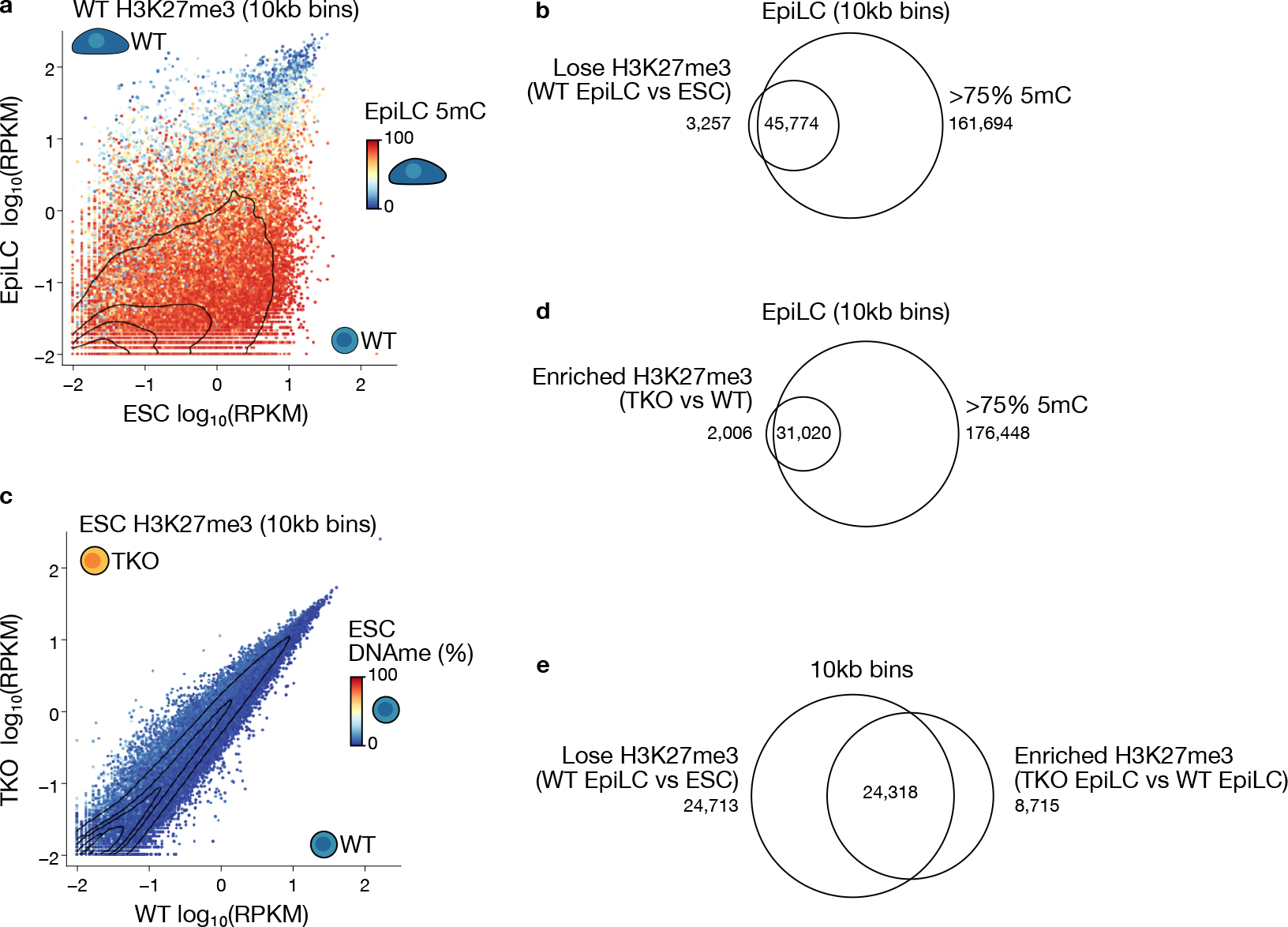
Genome-wide H3K27me3 dynamics during the exit from naïve pluripotency. **a.** 2D scatterplot showing H3K27me3 enrichment over genome-wide 10kb bins in differentiating WT cells. Datapoints are coloured by 5mC levels in WT EpiLCs. 10kb bins with ≥10 CpGs covered by ≥5 reads are shown (n=252,559), and the density of datapoints is included. **b.** Venn diagram showing the overlap between 10kb genomic bins that significantly (linear modelling using Limma, fold-change >2, t-test adjusted p <0.05) lose H3K27me3 in WT EpiLCs compared to WT ESCs and bins that are hypermethylated (>75% 5mC) in WT EpiLCs. **c.** 2D scatterplot showing H3K27me3 enrichment over genome-wide 10kb bins in WT and TKO ESCs. Data points are coloured by 5mC levels in WT ESCs. 10kb bins with ≥10 CpGs covered by ≥5 reads are shown (n=252,812), and the density of datapoints is included. **d.** Venn diagram illustrating the overlap between 10kb genomic bins with significant H3K27me3 level enrichment in TKO EpiLCs (vs WT EpiLCs) and bins that are normally hypermethylated (>75% 5mC) in WT EpiLCs. **e.** Venn diagram showing the overlap of 10kb bins that significantly lose H3K27me3 levels during differentiation from naïve ESCs to EpiLCs, and 10kb bins with aberrant enrichment of H3K27me3 levels in TKO EpiLCs compared to WT EpiLCs.

**Figure S4.**
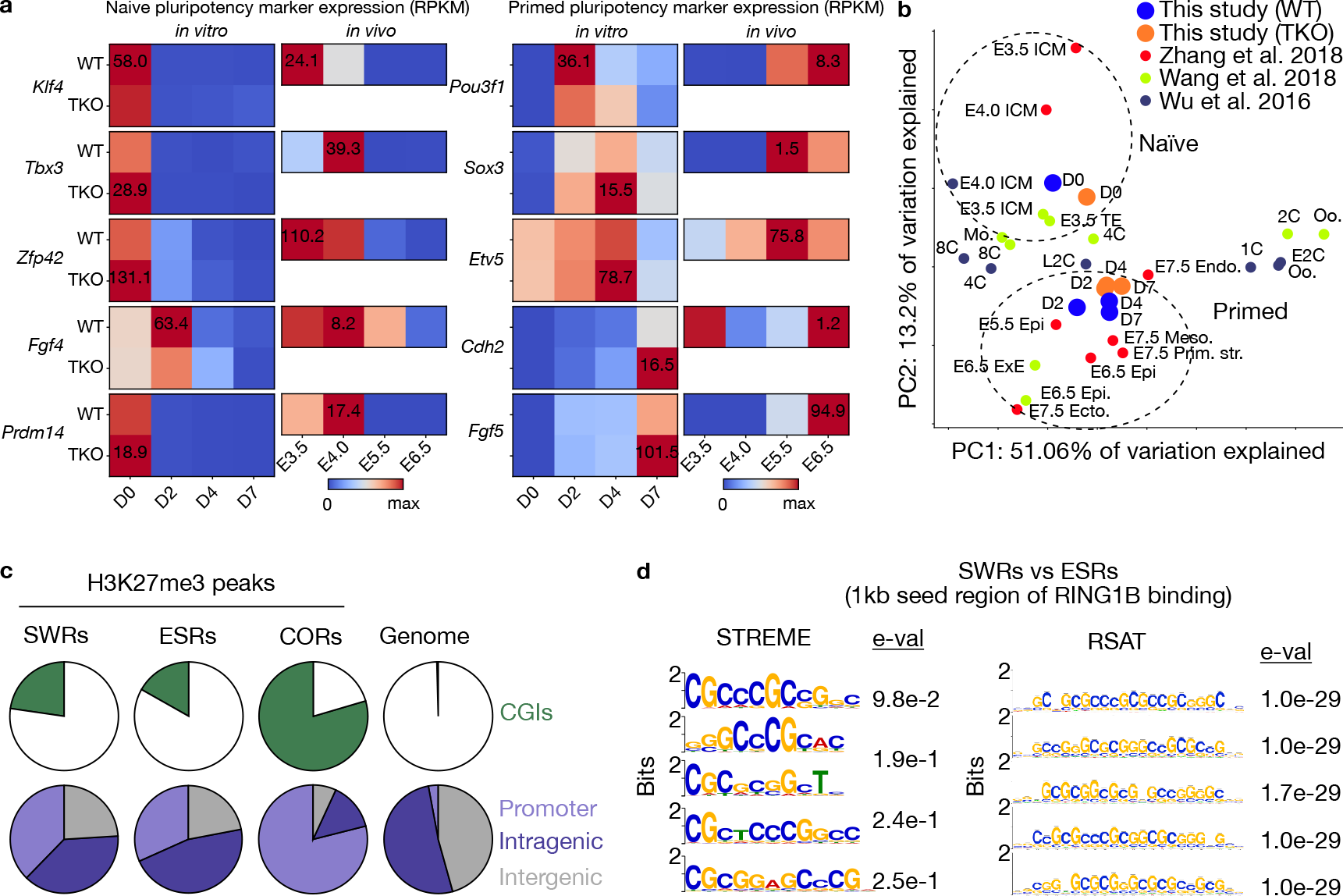
The transcriptomic landscape of TKO EpiLCs and genomic composition of SWRs. **a.** Heatmaps showing expression levels (RPKM) of naïve (*Klf4, Tbx3, Zfp42/Rex1, Fgf4 and Prdm14*) and formative/primed (*Pou3f1, Sox3, Etv5, Cdh2 and Fgf5*) pluripotency markers *in vitro* (0, 2, 4 and 7 days post-AF treatment) and *in vivo* (inner cell mass cells of E3.5 and E4.0 blastocysts, and E5.5 and E6.5 epiblasts). For each *in vitro* and *in vivo* dataset, the scale bar for each gene is set independently to the maximum expression of that gene in WT and TKO samples. **b.** Principal Component Analysis (PCA) plot showing transcriptomic similarities between naïve ESCs with the inner cell mass cells of the blastocyst, and between EpiLCs with epiblasts. Both groups are arbitrarily circled by a dotted line. D0-,2-,4-,7-: embryonic stem cells days post-AF treatment, Oo.: oocyte, 1C: zygote, E2C: early 2-cell embryo, 2C: 2-cell embryo, L2C: late 2-cell embryo, 4C: 4-cell embryo, 8C: 8-cell embryo, Mo.: morula-stage embryo, ICM: inner cell mass cells of the blastocyst, TE: trophectoderm cells of the blastocyst, E5.5-, 6.5-, 7.5-: embryonic days, Epi: epiblast, ExE: extraembryonic ectoderm, Endo.: endoderm, Meso.: mesoderm, Ecto.: ectoderm, Prim. Str.: primitive streak. **c.** Pie charts showing the genomic distribution of H3K27me3 peak classes <u>SW</u>itch <u>R</u>egions (SWRs), <u>E</u>SC-<u>S</u>pecific <u>R</u>egions (ESRs) and COnstitutive Regions (CORs). A fourth pie chart showing the total area of each genomic compartment is included. **d.** Motif analysis showing enriched motifs in the putative PRC binding sites within SWRs using ESRs as control sequences and two motif analysis algorithms.

**Figure S5.**
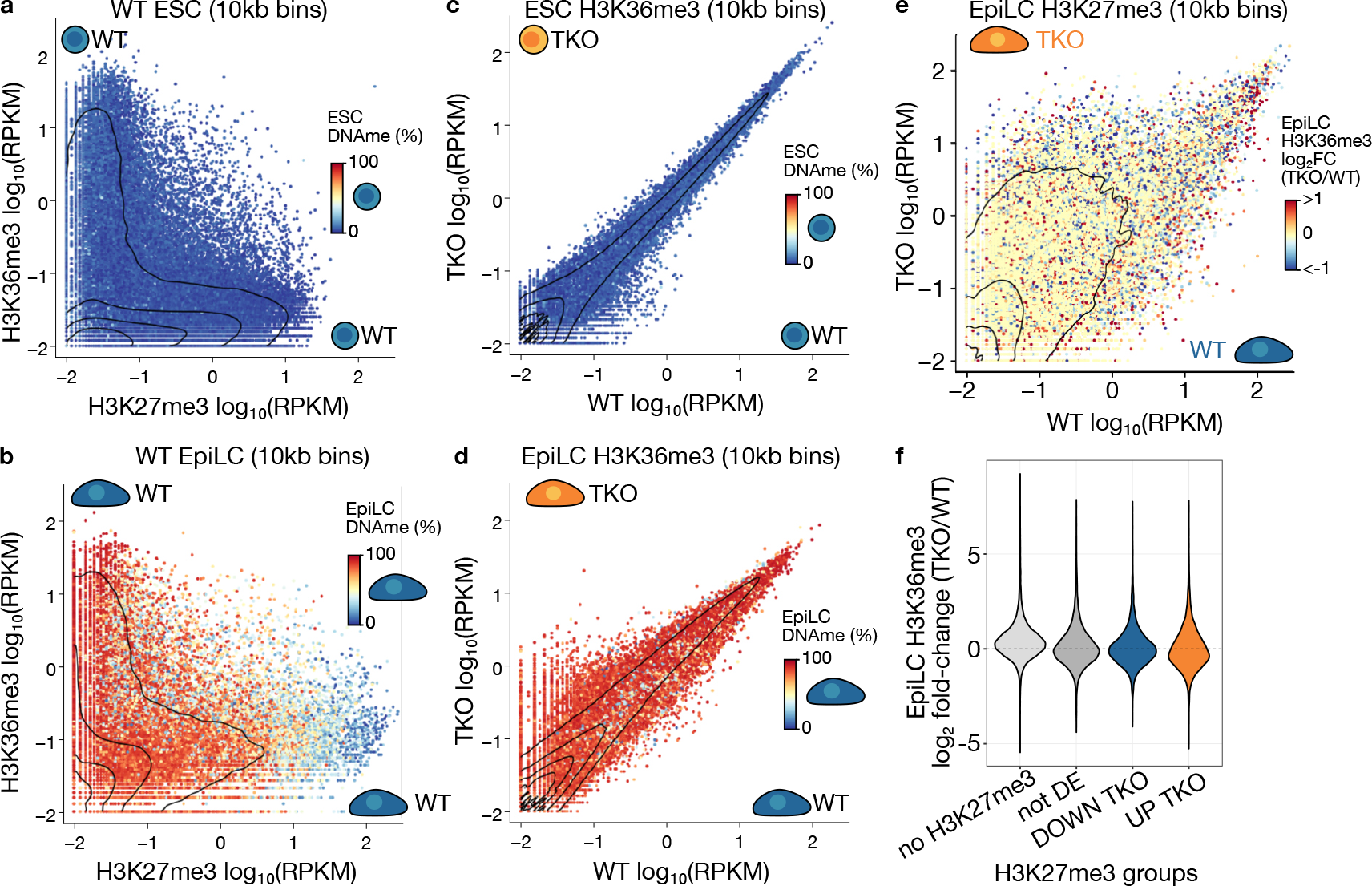
H3K36me3 deposition is not strongly impacted in absence of DNA methylation. **a.** 2D scatterplot showing H3K27me3 and H3K36me3 enrichment over genome-wide 10kb bins in WT ESCs. Data points are coloured by 5mC levels in WT ESCs. 10kb bins with ≥10 CpGs covered by ≥5 reads are shown (n=252,812), and the density of datapoints is included. **b.** 2D scatterplot showing H3K27me3 and H3K36me3 enrichment over genome-wide 10kb bins in WT EpiLCs. Data points are coloured by 5mC levels in WT EpiLCs. 10kb bins with ≥10 CpGs covered by ≥5 reads are shown (n=252,559), and the density of datapoints is included. **c.** 2D scatterplot showing H3K36me3 enrichment over genome-wide 10kb bins in WT and TKO ESCs. Data points are coloured by 5mC levels in WT ESCs as in **a**, and density of datapoints is included. **d.** 2D scatterplot showing H3K36me3 enrichment over genome-wide 10kb bins in WT and TKO EpiLCs. Data points are coloured by 5mC levels in WT EpiLCs as in **b**, and density of data points is included. **e.** 2D scatterplot showing H3K27me3 enrichment over genome-wide 10kb bins in WT and TKO EpiLCs, as in **Fig. 1c**. Data points are coloured by the relative change in H3K36me3 enrichment in TKO versus WT EpiLCs, and density of datapoints is included. **f.** Violin plot showing the distribution of H3K36me3 level change in TKO EpiLCs compared to WT EpiLCs. 10kb bins are categorized based on H3K27me3 dynamics in WT and TKO EpiLCs; No H3K27me3: insufficient H3K27me3 levels (CPM<1) in either WT or TKO EpiLCs, not DE: H3K27me3 levels do not significantly change between WT and TKO EpiLCs, Down TKO: relative loss of H3K27me3 in TKO EpiLCs, Up TKO: relative enrichment of H3K27me3 in TKO EpiLCs compared to WT EpiLCs.

**Figure S6.**
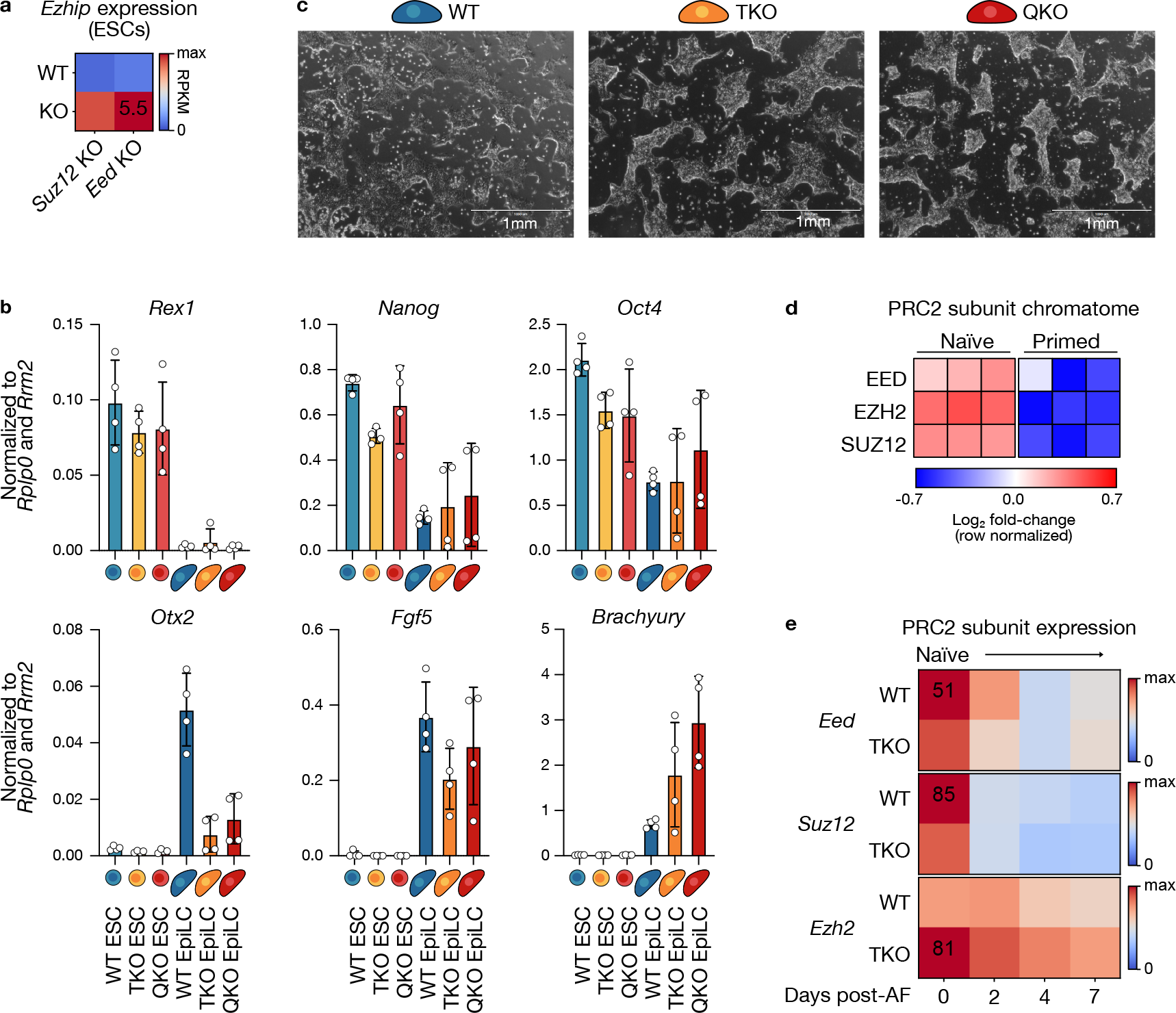
TKO+*Ezhip* KO (QKO) ESCs differentiate to EpiLCs. **a.** Heatmap showing *Ezhip* expression levels (RPKM) in WT and *Suz12* KO and *Eed* KO ESCs. The value of the highest expression level in all four conditions is indicated. **b.** Bar plots showing the expression level of naïve (*Rex1/Zfp42, Nanog*), general (*Oct4*), formative, (*Otx2, Fgf5*) and primed (*Brachyury/T*) pluripotency markers. Bars represent the mean, error bars the standard deviation, and individual data points are shown for each replicate. EpiLCs were collected 7 days post-AF treatment. **c.** Representative brightfield images of EpiLCs. Scale bar = 1mm. Pictures were taken 5 days post-AF treatment. **d.** Heatmap showing relative enrichment of PRC2 subunit proteins on the chromatin of WT naïve and primed epiblast stem cells (EpiSCs) (from Ugur *et al*., 2023). **e.** Heatmap depicting PRC2 subunit gene expression levels (RPKM) during EpiLC differentiation in WT and TKO cells.

**Figure S7.**
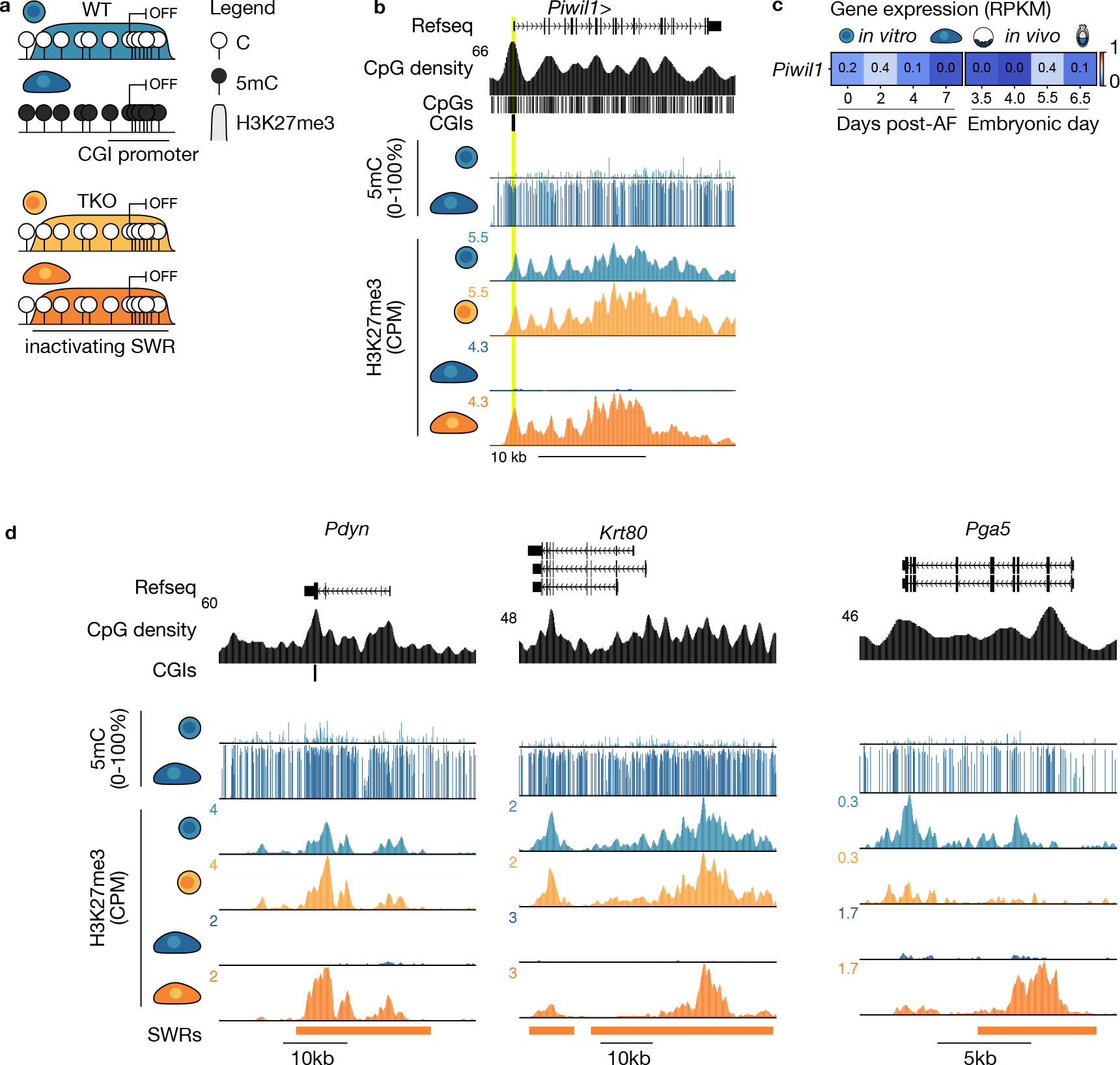
Examples of SWR regions with maintenance of gene repression or activation. **a.** Schema showing a hypothetical H3K27me3-to-5mC SWitch Region (SWR) that overlaps a genic promoter. In this case, dense methylation of the promoter region results in the maintained repression of expression. **b.** UCSC Genome Browser screenshot of an example SWR overlapping the *Piwil1* germline gene promoter. The Refseq gene annotation, CpG density, individual CpGs, CpG islands and scale bar are included. Note the anticorrelated enrichment of 5mC and H3K27me3 in WT cells, and the aberrant maintenance of H3K27me3 in TKO EpiLCs. Coordinates: chr5:128,733,977-128,756,805 **c.** Heatmap of *Piwil1* expression levels (RPKM) *in vitro* and *in vivo* differentiation. D0-,2-,4-,7-: embryonic stem cells days post-AF treatment. E3.5-, E4.5-: embryonic days 3.5 and 4.5 inner cell mass cells, E5.5-, E6.5: embryonic days 5.5 and 6.5 epiblast cells. **d.** UCSC Genome Browser screenshots of SWRs that overlap genes activated in WT EpiLCs (see schema, **Fig. 3b**). Coordinates: *Pdyn* chr2:129,673,296-129,713,102, *Krt80* chr15:101,345,735-101,394,892, *Pga5* chr19:10,666,677-10,680,350.

**Figure S8.**
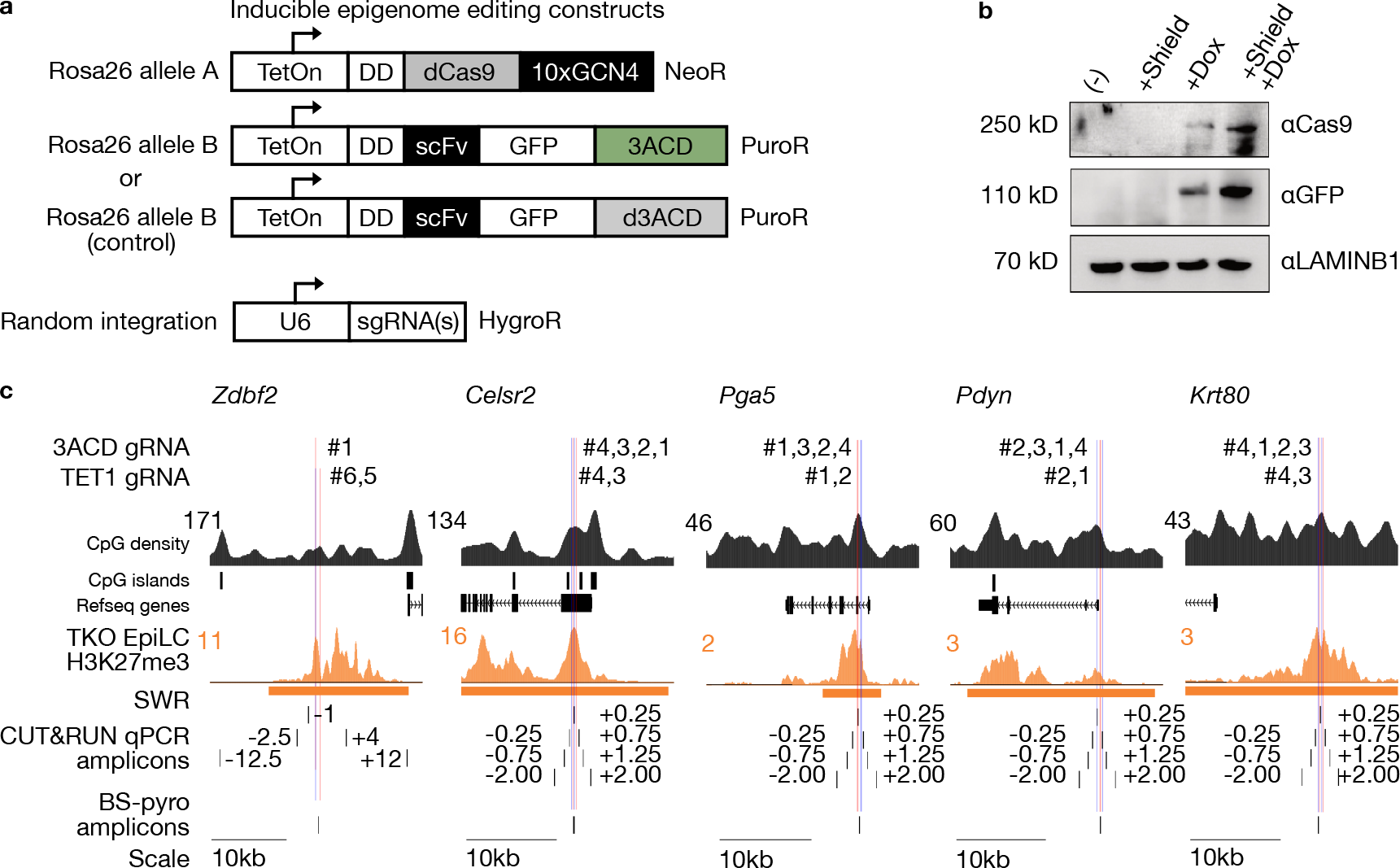
Implementation of an epigenome editing system for site-specific 5mC deposition at candidate SWRs. **a.** Schema showing inducible constructs integrated into the *Rosa26* locus in ESCs. One allele contains a construct composed of the TetOn array promoter for Doxycycline-dependent activation; DD: FKBP12-derived destabilizing domain, stabilized by the addition of the Shield-1 ligand; dCas9: catalytically dead Cas9, where two point mutations abrogate nucleolytic activity; 10xGCN4: an array of ten GCN4 epitopes (dCas9-SunTag). The homologous allele encodes a TetOn promoter, a destabilizing DD domain, a single-chain antibody (scFv) that recognizes the GCN4 epitope, green fluorescent protein (GFP) and the catalytic domain of DNMT3A (3ACD). Control cells contain an identical construct that encodes a catalytically inactive form of DNMT3A (d3ACD). A separate plasmid containing single guide RNAs (sgRNAs) was randomly integrated by piggyBac transposition. Resistance genes driven by an independent promoter are indicated. **b.** Western blot validating the inducible dCas9-SunTag/scFv-GFP-3ACD cell line. The two proteins (dCas9-SunTag and scFv-GFP-3ACD) encoded by the two inducible constructs are not detected in the absence of Dox, detected at intermediate levels after Dox induction, and robust expression after the addition of both Dox and Shield-1. **c.** UCSC Genome Browser screenshots of candidate SWR regions adjacent to the genes: *Zdbf2*, Celsr2, Pga5, Pdyn, and Krt80. The target site for 5mC editing using 3ACD and TET1 is shown. H3K27me3 CUT&Tag in TKO EpiLCs is included. The location of H3K27me3 CUT&RUN quantitative PCR and bisulphite-pyrosequencing (BS-pyro) amplicons are also shown. CpG density, CpG islands (CGIs) and Refseq gene annotations have been added for reference. Coordinates: *Zdbf2* chr1:63,246,946-63,275,173; *Celsr2* chr3:108,401,129-108,424,638; *Pga5* chr19:10,660,312-10,683,383; *Pdyn* chr2:129,683,316-129,707,017; *Krt80* chr15:101,366,622-101,390,127.

**Figure S9.**
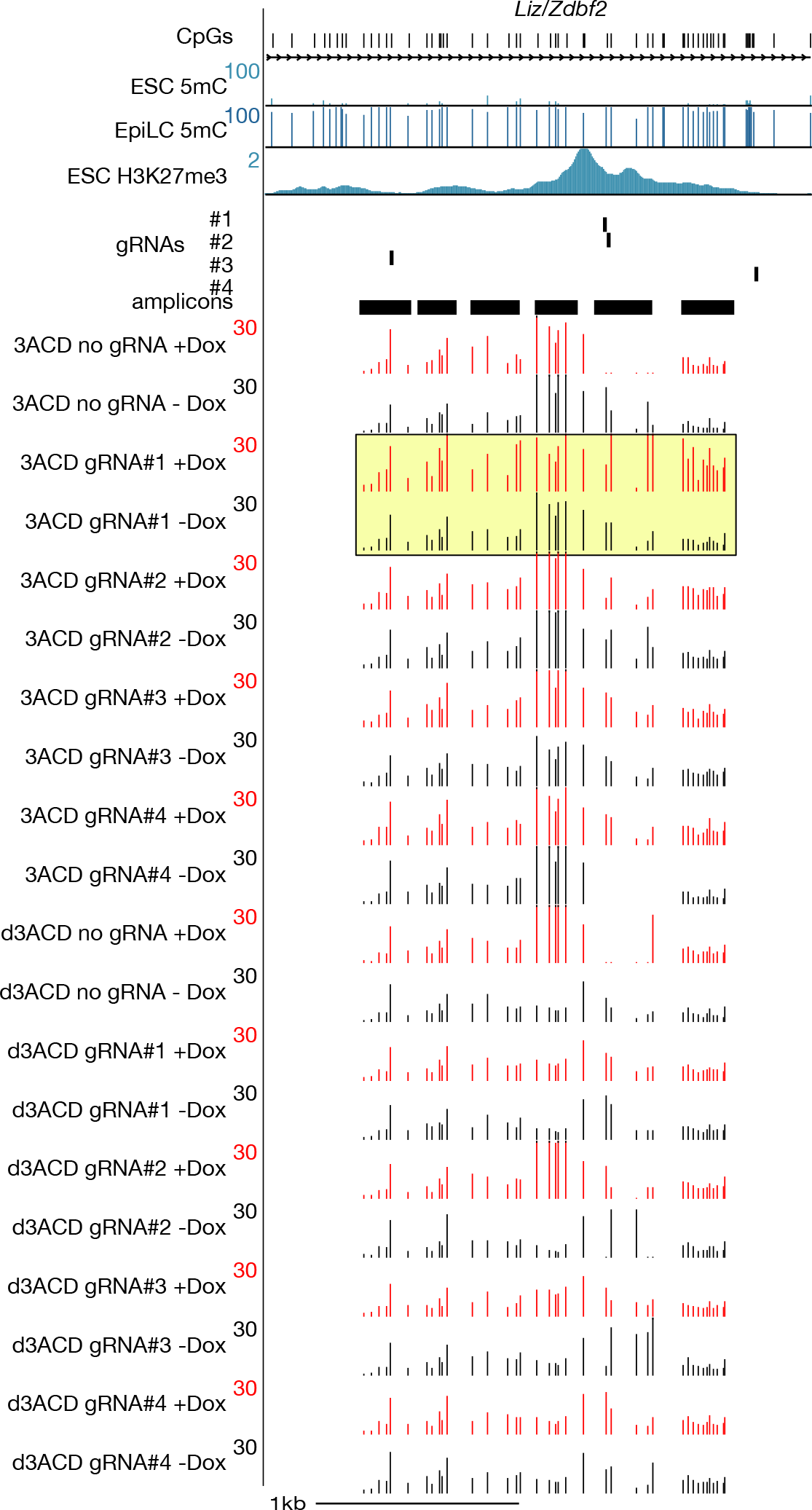
Site-directed 5mC induction at the *Zdbf2* SWR in ESCs. UCSC Genome Browser screenshot showing 5mC levels over six amplicons spanning the *Zdbf2* SWR. ESCs were grown in the presence (+Dox) and absence (-Dox) of Doxycycline, and four guide RNAs were tested in 3ACD and d3ACD cell lines. A no-gRNA control was also included (20 samples total). The genomic location of CpGs, Refseq annotations, WGBS from ESCs and EpiLCs, and H3K27me3 CUT&Tag from WT ESCs are shown. Coordinates: chr1:63,259,305-63,261,982.

**Figure S10.**
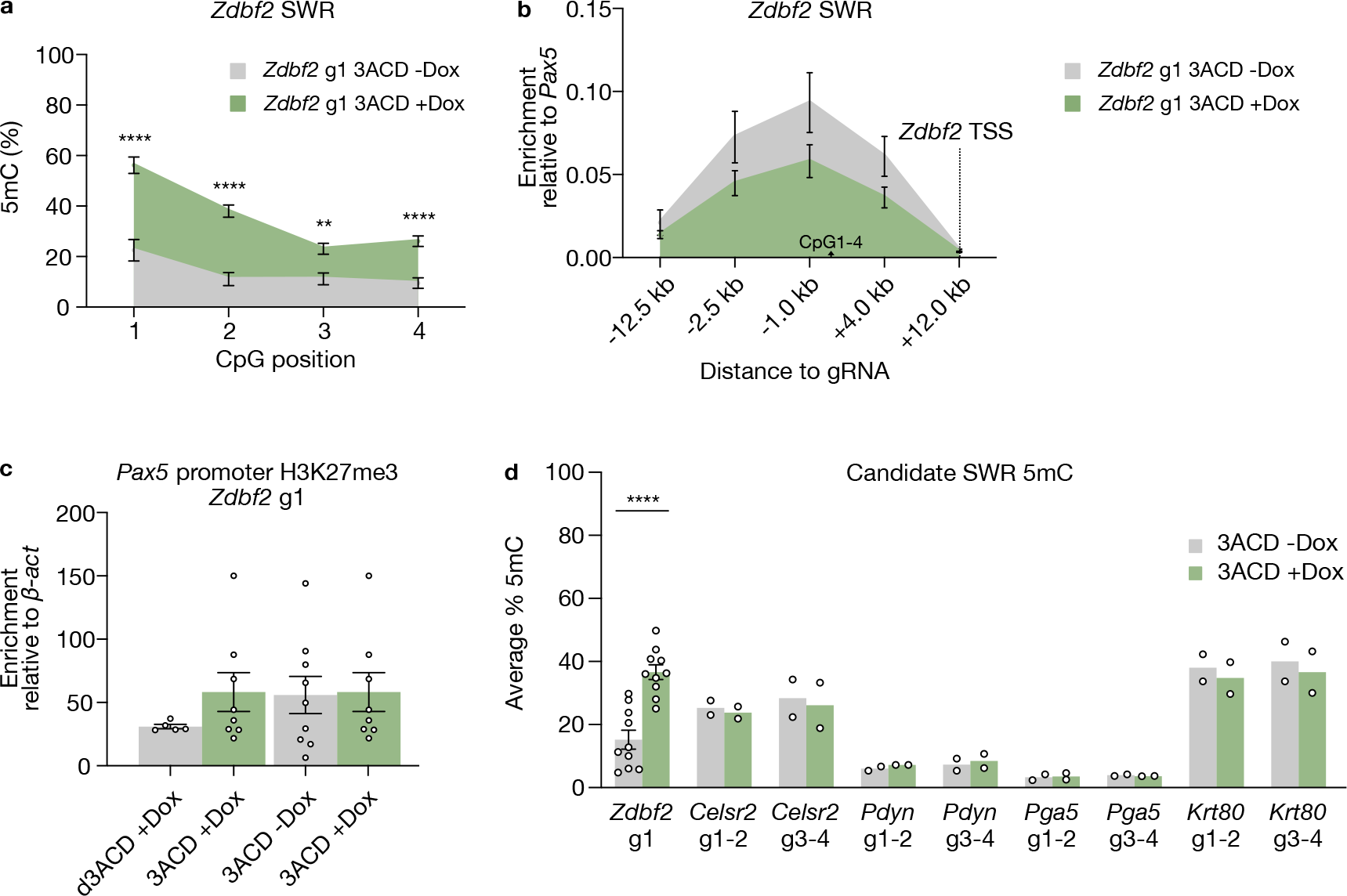
Targeted 5mC deposition and concomitant H3K27me3 loss at the *Zdbf2* SWR and site-directed epigenome editing at other candidate SWRs. **a.** Bisulphite-pyrosequencing of the *Zdbf2* SWR showing 5mC deposition in induced cells (+Dox +Shield-1, green) expressing dCas9 and the 3ACD construct compared to uninduced control cells (-Dox -Shield-1, gray). Cells were collected 7 days after the addition of Dox and Shield-1. Data represents mean of 9 replicates. **b.** CUT&RUN-qPCR showing lower average levels of H3K27me3 in induced cells (green) over the *Zdbf2* SWR. Data represent mean of 8 and 9 replicates for induced and uninduced samples, respectively. **c.** CUT&RUN-qPCR showing similar levels of H3K27me3 enrichment at the positive control *Pax5* promoter locus relative to a negative (background) control, *β-actin*. Data represents mean of 5-9 replicates, with each replicate indicated as a dot. **d.** Bisulphite-pyrosequencing results at the *Zdbf2, Celsr2, Pdyn, Pga5* and *Krt80* candidate SWRs. Each data point represents the average 5mC level over the pyrosequencing amplicon (3-5 CpGs) for each replicate, and bars represent the mean of replicates. Data are shown as mean ± standard error (for *Zdbf2*), single replicates are indicated as dots. p-values were calculated by two-tailed unpaired t-test assuming equal variance: **p<0.01, ****p<0.0001.

**Figure S11.**
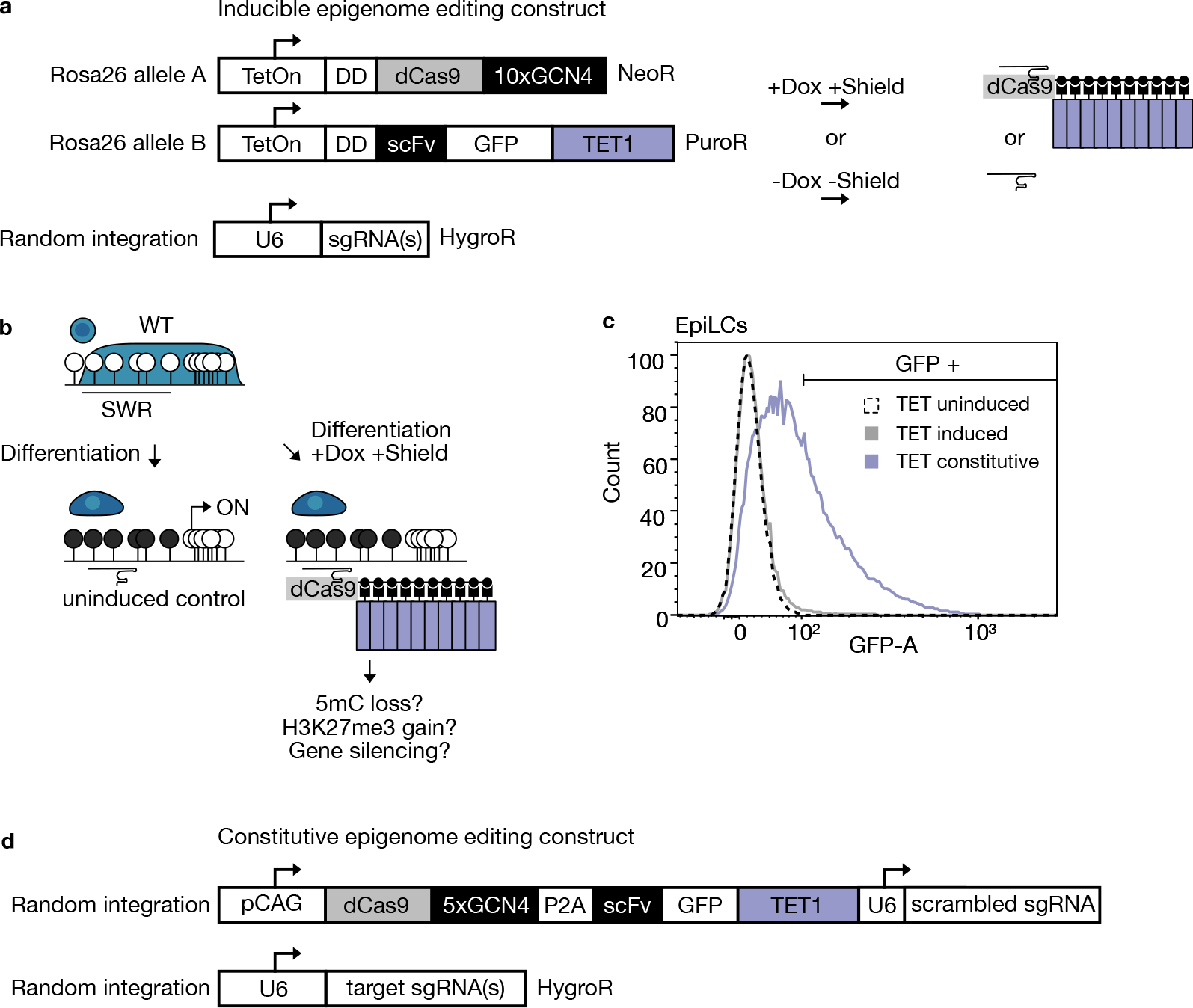
Targeted 5mC demethylation in inducible and constitutive systems. **a.** Schema showing inducible constructs integrated into the *Rosa26* locus in ESCs as in Figure S8. In this case, the mouse TET1 catalytic domain (TET1) is used, and uninduced (-Dox, -Shield) cells are used as controls. **b.** Schema of site-directed 5mC erasure at candidate SWRs using an inducible TET1-targetting system. Demethylation occurs after 5mC oxidation by the catalytic domain of the TET1 catalytic domain (purple). Site-specific demethylation is achieved by inducible expression of dCas9-SunTag/ scFv-TET1 and gRNAs complementary to candidate SWRs. Uninduced cells (-Dox -Shield) that only express the gRNA are used as a control. **c.** Histogram showing the distribution of GFP intensity in cells by flow cytometry. One representative experiment from three cell lines is shown for EpiLCs (4 days post-AF treatment): uninduced (-Dox -Shield, dotted line), induced (+Dox +Shield, grey) and constitutive TET1 (purple, see below) expression. **d.** Schema showing the constitutive epigenome editing strategy using a pCAG promoter and an all-in-one construct. Here, the SunTag is composed of 5 GCN4 epitopes, and is separated from scFv-GFP-human TET1CD by a P2A self-cleavable peptide. The construct was randomly integrated by piggyBac transposase, and expressing cells were selected using FACS.

**Figure S12.**
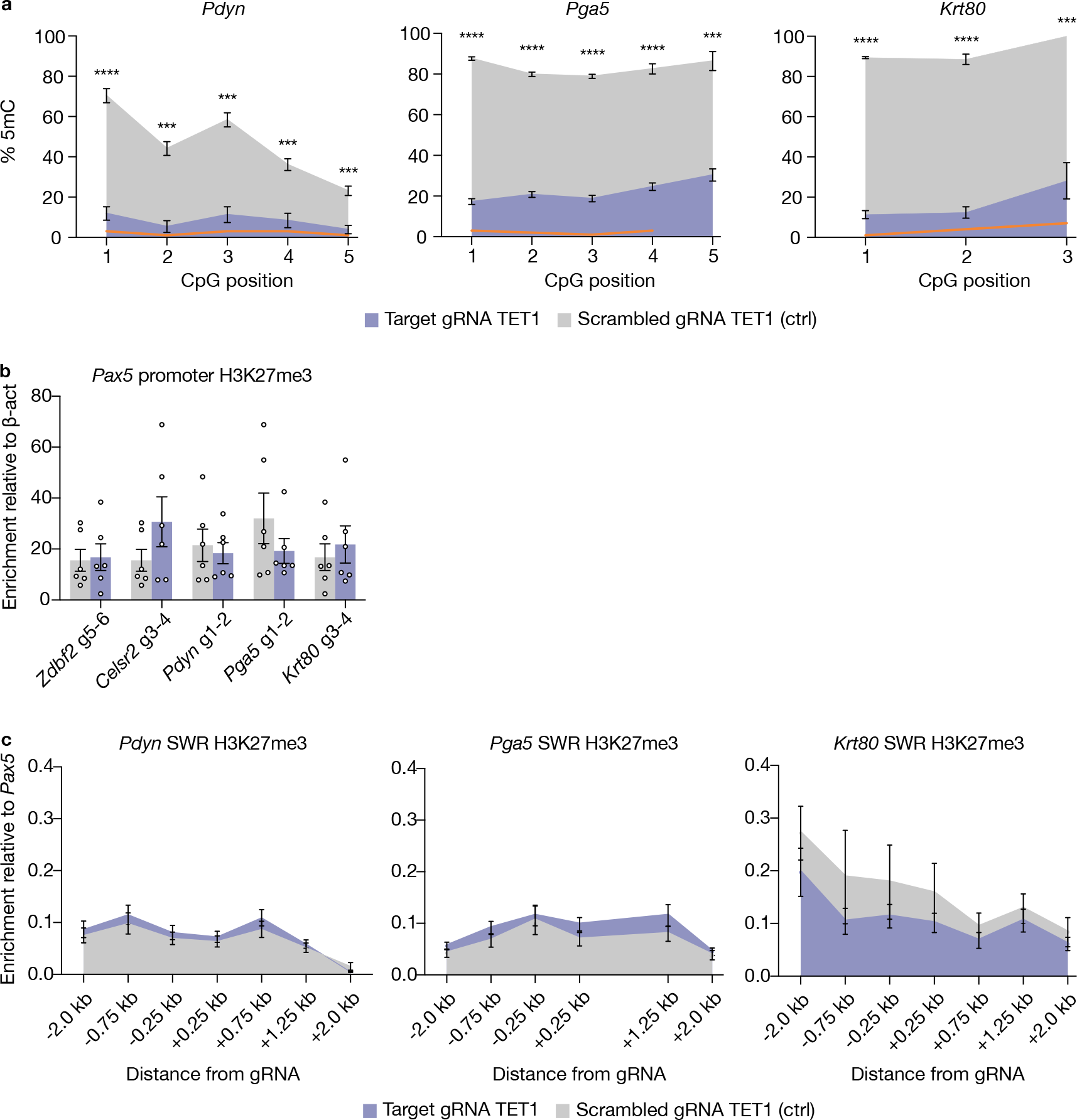
Targeted 5mC demethylation of candidate SWRs using the dCas9-SunTag/scFv-GFP-TET constitutive system. **a.** Bisulphite-pyrosequencing results from EpiLCs (4 days post-AF treatment) at candidate SWRs *Pdyn* (gRNA 1-2), *Pga5* (gRNA 1-2) and *Krt80* (gRNA 3-4). All cells constitutively express the dCas9-SunTag/ scFv-GFP-TET1CD construct. Cells expressing target gRNAs are in purple, and control cells expressing scrambled gRNAs are in grey. TKO EpiLCs were used as a negative control (orange line, 1 replicate). Mean 5mC levels at individual CpGs are shown. Data represent 4 replicates. **b.** CUT&RUN-qPCR results showing similar enrichment of H3K27me3 at the positive control *Pax5* locus across cell lines relative to *β-actin* (negative control locus). Data represent mean of 6 replicates, with replicates indicated as dots. **c.** H3K27me3 CUT&RUN-qPCR results shown as relative enrichment to *Pax5* (positive control locus) (ΔCt method). The amplicons analyzed span a region of 4 kb centered at the gRNA target regions. Error bars indicate ± standard error. p-values were calculated by two-tailed unpaired t-test assuming equal variance: \**P<0*.*05*, \*\**P<0*.*01*, \*\*\**P<0*.*001*, \*\*\*\**P<0*.*0001*.

**Figure S13.**
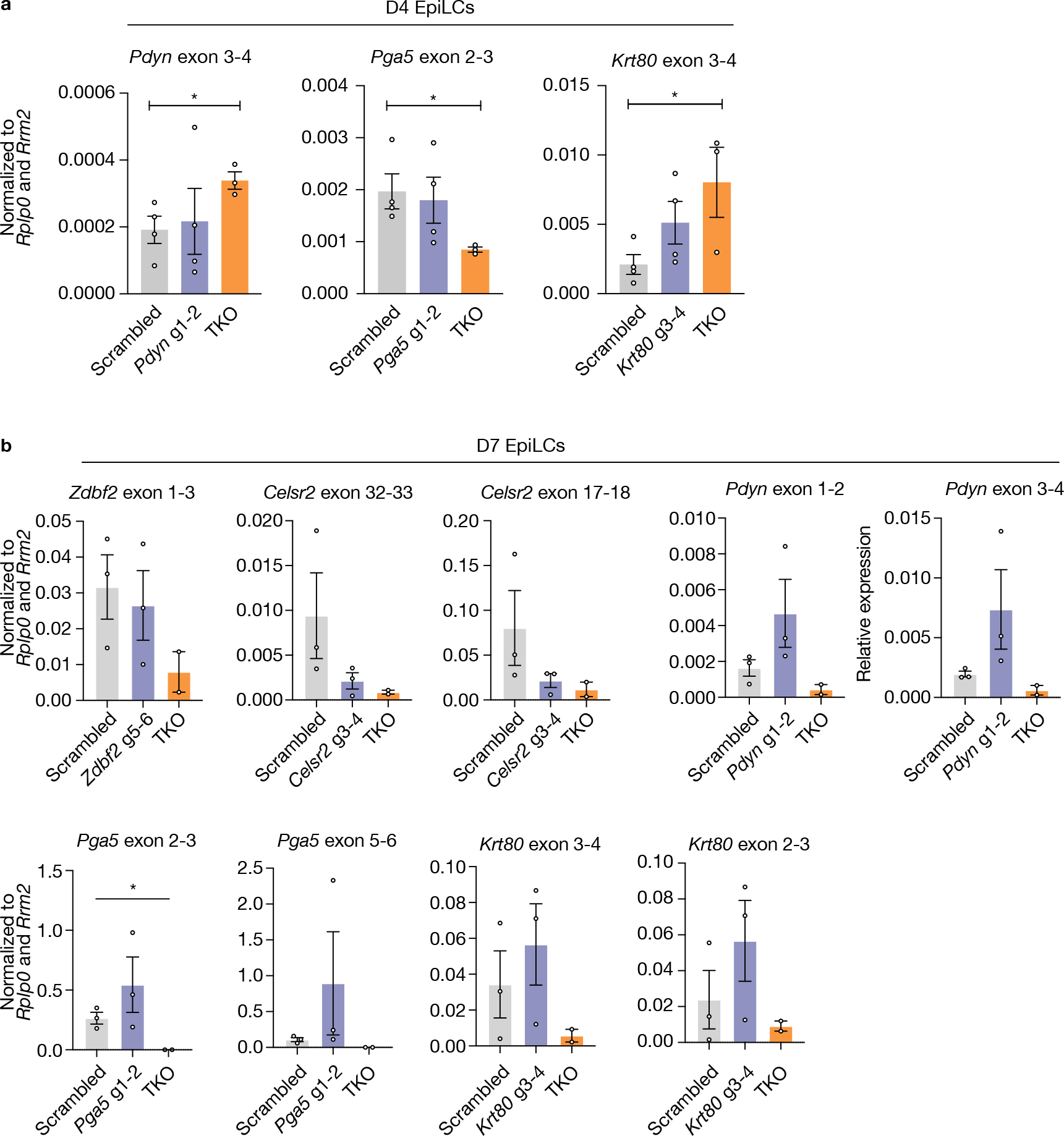
Expression analysis of genes adjacent to targeted 5mC demethylation loci in EpiLCs. **a.** RT-qPCR results for *Pdyn* exon 3-4, *Pga5* exon 2-3, and Krt80 exon 3-4 are shown as relative enrichment to geometric mean of two housekeeping genes (*Rplp0* and *Rrm2*) (ΔCt method). TKO EpiLC (4 days post-AF treatment) relative expression levels are also displayed. Data represent mean of 3-4 replicates, with replicates indicated as dots. **b.** RT-qPCR results from EpiLCs 7 days post-AF treatment are shown. Data presented as in **a**. Error bars indicate ± standard error. p-values were calculated by two-tailed unpaired t-test assuming equal variance. \**P<0*.*05*, \*\**P<0*.*01*, \*\*\**P<0*.*001*, \*\*\*\**P<0*.*0001*.

**Figure S14.**
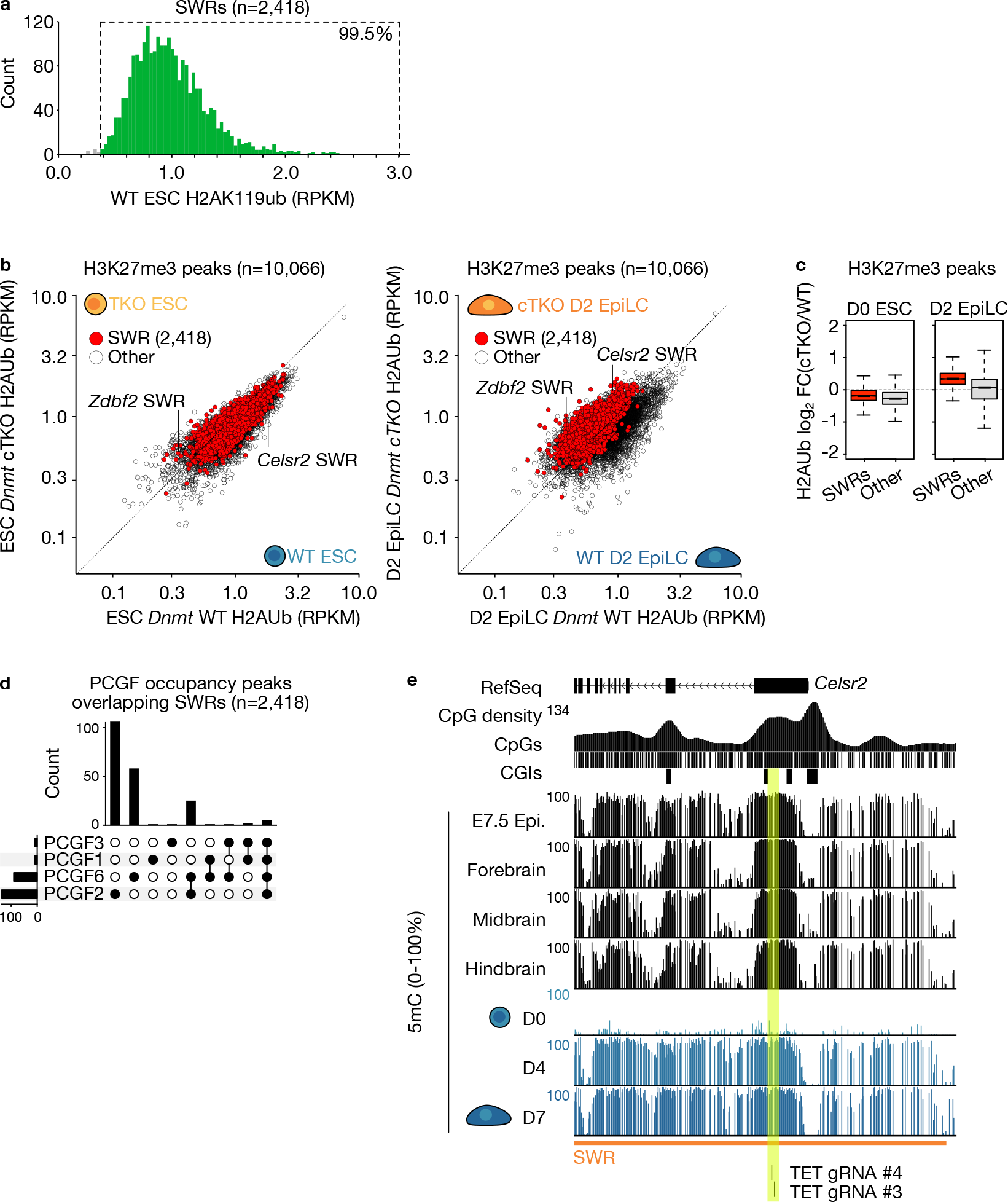
PRC1 and H2AK119ub analysis at SWRs. **a.** Histogram showing the distribution of H2AK119ub enrichment levels (RPKM) over SWRs (n=2,418) in hypomethylated ESCs. SWRs enriched for H2AK119ub (RPKM ≥0.5) are shown in green. **b.** 2D scatterplots showing H2AK119ub enrichment levels (RPKM) over H3K27me3 peaks (n=10,066) in WT and conditional TKO (cTKO) ESCs (left) and D2 EpiLCs (right, two days post-AF treatment). **c.** Boxplots showing the distribution of H2AK119ub level fold-change over H3K27me3 peaks in cTKO versus WT ESCs and D2 EpiLCs. **d.** UpSet plot showing the number of PCGF3, PCGF1, PCGF6 and PCGF2 peaks that overlap SWRs. **e.** WGBS tracks from various *in vivo* tissues where the *Celsr2* gene is expressed, indicating high 5mC content in the SWR region. The gRNAs used in this study are highlighted.

